# Mg^2+^-Dependent Multistep Folding and Stabilization of the GAAA Tetraloop-Receptor Interaction in a Group I Intron

**DOI:** 10.64898/2026.01.21.700762

**Authors:** Sk Habibullah, Dibyendu Mondal, Sunil Kumar, Govardhan Reddy

## Abstract

Group I Introns are non-coding regions of pre-mRNA that catalyze their splicing from the RNA sequence by folding to a specific structure. We used computer simulations to study the folding mechanism of the P4-P6 domain in the *Tetrahymena thermophila* group I intron, focusing on the GAAA tetraloop-receptor (TL-R) interaction, which is a ubiquitous tertiary interaction in RNA structures. We show that the intron folds via a multistep pathway, populating seven states with distinct tertiary contacts. Under physiological Mg^2+^ concentrations ([Mg^2+^]), the loop-bulge-P4 tertiary interaction is essential to stabilize the docked TL-R complex, whereas in high [Mg^2+^], the TL-R complex is stable by itself. The solvated Mg^2+^ ions modulate the TL-R docking–undocking dynamics and stabilize non-native intermediate states. The condensation of Mg^2+^ in the major grooves of the TL and R helices is critical for them to attain specific stiffness essential for their facile docking. The results highlight the critical role of Mg^2+^ ions in facilitating TL-R interaction formation, which stabilizes long-range tertiary contacts in RNA structures.

**For Table of Contents Use Only:** 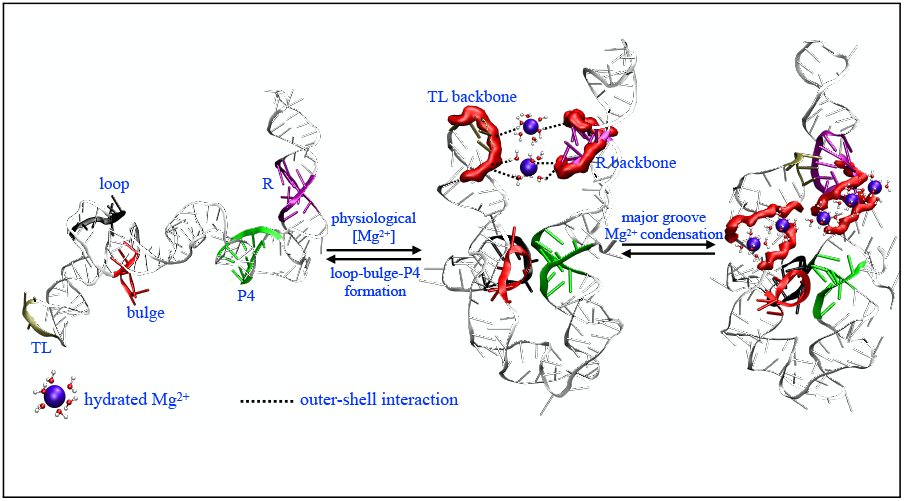

## Introduction

In the human body, there are approximately 20,000-25,000 protein-coding genes, yet it is estimated that more than 90,000 distinct proteins are produced to support essential biological functions. ^1–4^ This remarkable diversity arises because a single gene can give rise to multiple protein variants. Introns are non-coding regions within pre-mRNA that enable this process by facilitating alternative splicing ^5,6^ in eukaryotic cells, thereby contributing to the generation of diverse mRNA transcripts and proteins.^7,8^ These non-coding regions of RNA play key roles in genome regulation, gene expression control, and RNA processing. ^9^ Additionally, introns act as sources for functional non-coding RNAs, such as microRNAs and snoRNAs, that further influence gene regulation and RNA modification.^10,11^

Group I introns are a distinct class of catalytic RNAs that self-splice via a two-step transesterification process^12,13^ and they are excellent models for studying RNA catalysis and structure-function relationships. ^14,15^ They are typically 250-500 nucleotides long and fold into conserved tertiary structures that enable protein-independent catalytic activity (Fig. 1A). ^16^ The active site in group I introns lies between the conserved regions (P4-P6 and P3-P9), allowing precise RNA cleavage and ligation (Fig. 1A,B).^17^ The discovery of group I introns was a major milestone in molecular biology, proving that RNA can function as an enzyme and broadening the known roles of RNA in biology.^18^

**Figure 1:**
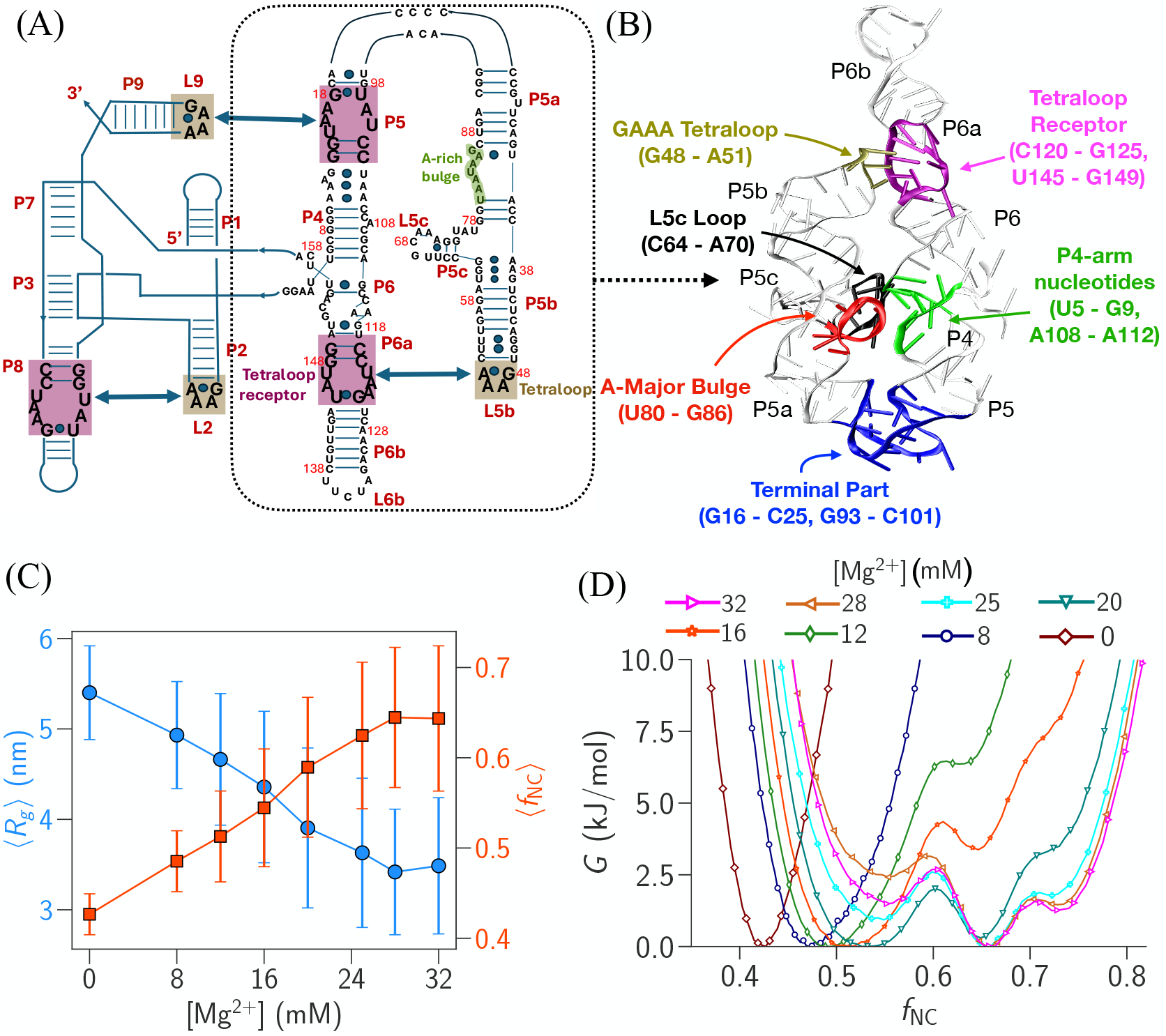
(A) Schematic of the secondary structure of group I intron from *Tetrahymena thermophila* showing GAAA tetraloop (olive) and receptor (purple) interactions. The Secondary structure of P4-P6 domain of the intron is highlighted using the dotted box. (B) Tertiary structure of P4-P6 domain of group I intron. Critical structural components GAAA tetraloop (olive), receptor (purple), L5c loop (black), A-major bulge (red), P4 helix nucleotides (green) and terminal part (blue) are shown. (C) Average fraction of the native contacts (⟨*f*_NC_⟩) and average radius of gyration ( ⟨*R*_*g*_⟩) of the P4-P6 domain of the intron are plotted as a function of [Mg^2+^]. (D) FES of the intron projected on *f*_NC_ at different [Mg^2+^].

Among the tertiary interactions present in the intron structure, tetraloop-receptor tertiary interactions play a critical role in the folding of group I introns in *Azoarcus* and *Tetrahymena* bacteria.^19–25^ These interaction acts as a structural anchor by bridging distant helices, thereby aligning and stabilizing the intron’s core architecture necessary for efficient self-splicing activity (Fig. 1A,B).^26,27^ The group I intron folding studies show that tetraloop-receptor interactions form early in the folding process and act as a scaffold for the formation of other tertiary interactions present in the structure.^19,20,23^ Depending on their location in the structure, early formation of these interactions, which bridge the distant helices, could also lead to the formation of topological traps, which slow down intron folding as the RNA has to escape from these traps to fold to the functional structure. ^25^

The GAAA tetraloop–receptor (TL-R) complex is an conserved RNA tertiary motif that is critical to the folding and stability of many large structured RNAs, including introns, riboswitches, ribosomal RNAs, and Ribonuclease P.^26,28–34^ As a member of the GNRA tetraloop (N=any nucleotide, R=A or G) family,^35^ the GAAA tetraloop adopts a distinctive U-turn conformation that exposes specific bases for interactions with distant receptor elements. This configuration enables the formation of specific hydrogen bonds and stacking interactions − known as A-minor interactions − with an 11-nucleotide receptor.^36,37^ The exceptional stability and modularity of the TL-R interaction make it an effective thermodynamic clamp promoting long-range tertiary contacts that anchor and organize complex RNA architectures.^27^ Due to the robust interaction patterns in the TL-R complexes, it is also used as a foundational module in RNA engineering and nanotechnology.^38,39^ Hence, it is essential to study the mechanism of TL-R interaction and the role of metal ions in the interaction.

The P4-P6 domain in the *Tetrahymena thermophila* group I intron is extensively used as a model system to understand the mechanism of TL-R interaction (Fig. 1A,B). The single-molecule fluorescence resonant energy transfer (smFRET) experiment studies on this system have shown that the folding dynamics, especially the docking and undocking processes, are highly dependent on the type and concentration of metal cations in the solution. ^40,41^ Studies proposed that the facilitation of TL-R docking by Mg^2+^ ions arises predominantly from entropic effects. ^42^ Higher Mg^2+^ concentration ([Mg^2+^]) did not significantly affect the enthalpy of docking, leading to the proposal that higher [Mg^2+^] reduced the entropy cost of counterion localization and receptor ordering. ^42^ Experiments also showed that the P4-P6 domain of the intron folds cooperatively, and condensation of the Mg^2+^ ions around the loop-bulge-P4 region (Fig. 1B) triggers compaction to native tertiary structure. ^43^ Theoretical analysis using the virtual bond RNA folding model (Vfold) and tightly bound ion model (TBI) models revealed that Mg^2+^ ions promote TL-R docking primarily through entropic stabilization. Mg^2+^ ions condense in the TL-R region and replace the Na^+^ ions, which are released into the solution, increasing the entropy of the solution. ^44^ Numerous simulation studies have probed the mechanism of isolated tetraloop formation and have highlighted their importance in RNA folding and function. ^45–54^ However, it is still not fully understood how Mg^2+^ ions directly facilitate TL-R complex formation and how other tertiary interactions support this process, which collectively governs the overall folding landscape of the RNA.

Experiments have shown that at optimal Mg^2+^ concentrations, the *Tetrahymena thermophila* group I intron folds rapidly and efficiently, ^55,56^ as Mg^2+^ stabilizes the native structure while preventing kinetic traps. ^21^ Incorrect tertiary interactions in the intron can result in a stable misfolded trap. In contrast, timely and accurate formation of specific tertiary contacts guides folding along productive pathways, reducing misfolding and accelerating attainment of the functional structure of intron. ^57,58^ Small angle X-ray scattering (SAXS) experiments on the P4-P6 domain of the *Tetrahymena thermophila* intron showed that low [Na^+^] favors an extended state without significant tertiary interactions. At [Na^+^] = 100 mM, a relaxed state with partial tertiary contacts without TL-R complex formation is stabilized. Where as at high ion concentrations, [Na^+^] = 1000 mM or [Mg^2+^] = 1 mM, the fully folded state is stable. ^59^ Furthermore, the smFRET experiments on the P4-P6 domain of the *Tetrahymena thermophila* intron inferred the stepwise formation of the tertiary interactions in the presence of Mg^2+^ ions.^60–64^

In this work, we performed extensive simulations to complement the experimental studies and gain insight into the structure of the intermediates populated during the TL-R complex formation and the role of metal ions in stabilizing them. We initially performed CG simulations to map the folding landscape of the P4-P6 domain of the group I intron. We demonstrate that the P4-P6 domain of the group I intron folds through a multistep pathway, populating eight distinct states with different tertiary contacts. The TL-R docked complex is stabilized by loop-bulge-P4 tertiary interactions, and its stability increases with the Mg^2+^ concentration ([Mg^2+^]) (Fig. 1B). In high [Mg^2+^], the TL-R docked complex is stable even without the loop-bulge-P4 tertiary interactions. The metadynamics simulations using an all-atom RNA model showed that the solvated Mg^2+^ ions influence the docking-undocking dynamics of the TL-R complex. The TL-R complex can adopt intermediate states lacking native hydrogen bonds, but are stabilized by water-mediated outer-shell interactions between Mg^2+^ ions and RNA phosphate oxygens. Interestingly, the condensation of Mg^2+^ ions in the major grooves of the TL and R helices helps maintain the critical helical lengths required for TL-R docking. This study offers insights into the formation of TL-R interactions, which are ubiquitous tertiary interactions that facilitate long-range contacts in folded RNA structures.

## Materials and Methods

### Computer Simulations

#### Coarse-Grained (CG) MD Simulations

The CG simulations were carried out using the three interaction site (TIS) RNA model.^65,66^ In this model, each RNA nucleotide is represented as three distinct beads representing phosphate, sugar, and base groups. The detailed description of the TIS RNA model can be found in references.^65–68^

The native hydrogen bond network required for the native-centric TIS RNA model was extracted from the high-resolution crystal structure of the P4-P6 domain of the group I intron (PDB ID: 1GID)^69^ using RNApdbee 2.0 server.^70^ The hydrogen bonds are given in Table S1, S2, and S3. The RNA and ions parameters are given in Table S4.

All CG Langevin dynamics simulations were performed using the *LangevinIntegrator* module in the OpenMM simulation package.^71^ The simulations were performed in the low-friction regime (*γ* = 0.01 ps^−1^)^72^ for enhanced conformational sampling. Long-range Coulomb interactions were treated via the Particle-Mesh-Ewald (PME) method as implemented in OpenMM. All the simulations were performed in a cubic simulation box of edge length 20 nm with RNA and ions explicitly present in the simulation box. The water solvent was treated implicitly. The temperature (*T*) was maintained at 298 K and we used a 2 fs integration time step. To investigate the impact of Mg^2+^ concentration ([Mg^2+^]) on the conformational ensemble of the intron, we performed simulations under the following conditions: (i) [Mg^2+^] = 0, 8, 12, 16, 20, 25, 28, 32, 50, and 125 mM with [Na^+^] fixed at 150 mM. For each [Mg^2+^], the cumulative simulation time is ≈ 11 *µ*s. The initial 1 *µ*s of data from each trajectory was discarded for calculating thermodynamic properties. All-atom conformations were reconstructed from the CG conformations using the TIS2AA tool. ^73^ We used VMD for visualization and rendering structural representations.^74^

### All-Atom (AA) MD Simulations

To investigate the role of Mg^2+^ ions in the formation of the TL-R complex, we performed well-tempered metadynamics (WT-MetaD) simulation using PLUMED 2.5.1^75,76^ and GROMACS 2018.6 software.^77,78^

#### Unbiased MD Simulations

Initially, we performed unbiased molecular dynamics (MD) simulations to equilibrate the system. The equilibrated conformations were subsequently used to perform the WT-MetaD. We used the modified AMBER ff14 force field^79^ (DESRES ff) and GROMACS 2018.6 simulation software for the simulations. We removed the phosphate group from the 5^′^-end of the X-ray crystal structure of the P4-P6 domain of the group I intron (PDB ID: 1GID). ^69^ The simulations were performed in a cubic box of length 13.5 nm with periodic boundary conditions in all directions. We added Na^+^ ions to neutralize the net charge in the system, and the simulation box was solvated with TIP4P-EW^80^ water molecules.

We performed energy minimization of the system using the steepest descent algorithm with a maximum force cutoff of 1000 kJ/(mol · nm). The system was equilibrated in three consecutive steps: (i) NVT simulation of solvent molecules at temperature *T* = 298 K for 150 ns by restraining the positions of RNA atoms and ions using a harmonic potential with a force constant of 1000 kJ/(mol · nm^2^), (ii) NVT simulation of solvent molecules and ions at *T* = 298 K for 150 ns by restraining the positions of RNA atoms and (iii) NPT simulation of solvent molecules and ions at *T* = 298 K and pressure *P* = 1 atm for 150 ns by restraining the positions of RNA atoms.

To generate the undocked TL-R complex, we heated the system from 300 K to 350 K by gradually increasing the temperature in intervals of 10 K (Fig. S1). We extracted the undocked TL-R complex and rebuilt the simulation box as described previously to perform the unbiased simulations of the undocked TL-R complex at *T* = 300 K and *P* = 1 atm. We added 150 Na^+^ ions ([Na^+^] = 100 mM) and 15 Mg^2+^ ions ([Mg^2+^] = 10 mM) to the simulation box. The simulation box was net charge neutralized by adding the required number of Cl^−^ ions. The monovalent and divalent ion parameters are taken from ref.^81,82^ We followed the same minimization and equilibration protocols as described before. After equilibration, we performed NPT simulation at *T* = 300 K and *P* = 1 atm without any restraints on all the atoms. The *T* and *P* are maintained using velocity rescaling thermostat^83^ with a time constant *τ*_*t*_= 0.1 ps and Parrinello-Rahman barostat^84^ with a time constant *τ*_*p*_ = 2 ps. We truncated the Lennard-Jones interactions at 0.1 nm. The electrostatic interactions were calculated using the particle-mesh Ewald (PME)^85^ method with a grid size of 0.16 nm and real space cutoff of 0.1 nm. All the covalent bonds containing hydrogen atoms are kept rigid using the LINCS algorithm.^86^ The equation of motion is integrated using the leapfrog integrator with a time step of 2 fs.

#### Well-Tempered Metadynamics Simulations

We took the equilibrated undocked TL-R complex and performed well-tempered metadynamics simulations (WT-MetaD)^87,88^ using PLUMED 2.5.1^75,76^ and GROMACS 2018.6 to probe the effect of Mg^2+^ ions on the free energy landscape of TL-R complex. We chose two collective variables (CVs) to project the free energy: (i) center of mass distance (*d*_*com*_) between 4 nt TL (G48-A51) and 11 nt R (C120-G125 and U145-G149) helices, (ii) inter helix native contact^89^ 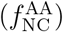 between 4 nt TL and 11 nt R helices where we used a total of 278 inter-helix distances (distances are between the heavy atoms) to calculate 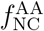 (Table S5).

We initiated the WT-MetaD simulations using the previously equilibrated systems from the unbiased MD simulations. We ran NPT simulations at *T* = 298 K and *P* = 1 atm for the TL-R system. The Gaussian widths for *d*_*com*_ and 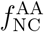 in the WT-MetaD simulations are 0.04 nm and 0.01 respectively. Initial Gaussian height, deposition time, and bias factor are 0.5 kJ/mol, 1 ps, and 15, respectively. For the CV, *d*_*com*_, we implemented a lower wall cut-off of 0.9 nm and an upper wall cut-off of 1.7 nm. We used sum hills and REWEIGHT BIAS modules embedded in PLUMED for reweighting simulation data. The module sum hills is used for projecting the free energy surfaces (FESs) onto the biasing CVs (*d*_*com*_ and 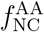). The module REWEIGHT BIAS is used to reweight the probability distributions and free energy surfaces onto other CVs. The convergence of WT-MetaD simulations was assessed by calculating the free energy profile as a function of three different CVs at various simulation times (Fig. S2A,B). Minimal changes in the FES with increasing time indicated convergence. This was further corroborated by the block analysis of the last 500 ns of the trajectory, which provided an estimate of the error in free energy projected onto *d*_*com*_ and 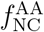 (Fig. S2C,D,E).

## Results and Discussions

### P4-P6 Domain in the Intron Populates Intermediates in its Folding Pathway

The physiologically abundant Mg^2+^ divalent cation plays a key role in intron folding because the RNA backbone is negatively charged.^90,91^ We studied the effect of Mg^2+^ concentration ([Mg^2+^]) on the folding thermodynamics of the P4-P6 Domain in the intron (from now on referred to as intron) using the CG TIS RNA model (see Methods) (Fig. 1B). The [Mg^2+^] was varied from 0 to 32 mM, while maintaining a fixed background concentration of Na^+^ at 150 mM. We computed ⟨*f*_NC_⟩ and ⟨*R*_*g*_⟩ of the intron to investigate the effect of [Mg^2+^] on its folding (Table S5, Fig. 1C). The plot shows that there is no sharp single transition in ⟨*f*_NC_⟩ or ⟨*R*_*g*_⟩ with increasing [Mg^2+^]. Instead, there is a gradual increase in ⟨*f*_NC_⟩ and a gradual decrease in ⟨*R*_*g*_⟩ with increasing [Mg^2+^], suggesting a progressive shift of the overall population toward the folded states. The simulation trajectories show that intron populates multiple distinct intermediates in its folding pathway (Fig. S3A), and with the increase in [Mg^2+^], the population shifts more towards the folded state.

We projected the free energy surface (FES) on *f*_NC_ to identify different thermodynamic states of the intron (Fig. 1D). For 0 < [Mg^2+^] < 12 mM, only the unfolded state (U) is populated at *f*_NC_ ≈ 0.43 − 0.49. However, with the increase in [Mg^2+^] from 0 to 12 mM, we observed a compaction of the unfolded state due to the decrease in the persistence length of the backbone from the screening of negative charge by Mg^2+^. As a result of the compaction, *f*_NC_ increased from 0.42 to 0.49 (Fig. 1D). For 16 < [Mg^2+^] < 32 mM, the intron populated two additional states which are classified as intermediate (I) and folded (F) states at *f*_NC_ ≈ 0.66 and ≈ 0.73, respectively (Fig. 1D). With an increase in [Mg^2+^], the stability of these two states increased.

For the intron to fold to its native state, three critical tertiary interactions need to be formed: (i) terminal region (G16-C25 and G93-C101), (ii) loop-bulge-P4 region (C64-A70, U80-G86, U5-G9 and A108-A112) and (iii) docking between TL and R to form the TL-R complex (G48-A51, C120-G125 and U145-G149) (Fig. 1B). Although the FES with *f*_NC_ as a CV shows three states (U, I, and F), it cannot resolve the stability of the three tertiary interactions in those three states. We quantified the extent of formation of the tertiary interactions by computing the fraction of native contacts corresponding to the terminal region 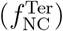, TL-R complex 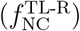, and loop-bulge-P4 region 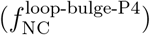 (Table S5). We studied the effect of [Mg^2+^] on the stability of these three tertiary interactions by computing the 1D FES for their formation projected onto their fraction of native contacts (Fig. S4A,B,C). As [Mg^2+^] increases, the basin with a higher fraction of native contacts is more populated in all three cases, suggesting that Mg^2+^ ions promote the formation of these tertiary contacts in the intron structure. We note that the F state (*f*_NC_ ≈ 0.73, 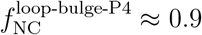) is not the global minimum in the FES even at high [Mg^2+^] (Fig. 1D, S4C). This is due to the drawback of the lack of explicit water molecules in the CG TIS model and it is discussed in the later subsections.

### Intron Folding Exhibits Kinetic Partitioning

We obtained the complete folding landscape of the intron from the CG simulations. The folding progresses through three major states: the unfolded state (U) ↔ the relaxed state (RL) ↔ the folded state (F) (Fig. 2 and S5). Using the three collective variables − (i) 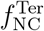, (ii) 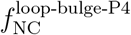, and (iii) 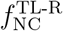− we constructed the folding landscape of the P4–P6 domain of the group I intron in the presence of Mg^2+^ (Fig. 3A,C). In the U state, all secondary contacts within the tetraloop and receptor helices are present, but the tertiary interactions are missing. The tertiary interactions emerge progressively in the RL and F ensembles. The RL and F states further subdivide into three substates each: R1, R2, R3, and F1, F2, F3, respectively.

**Figure 2:**
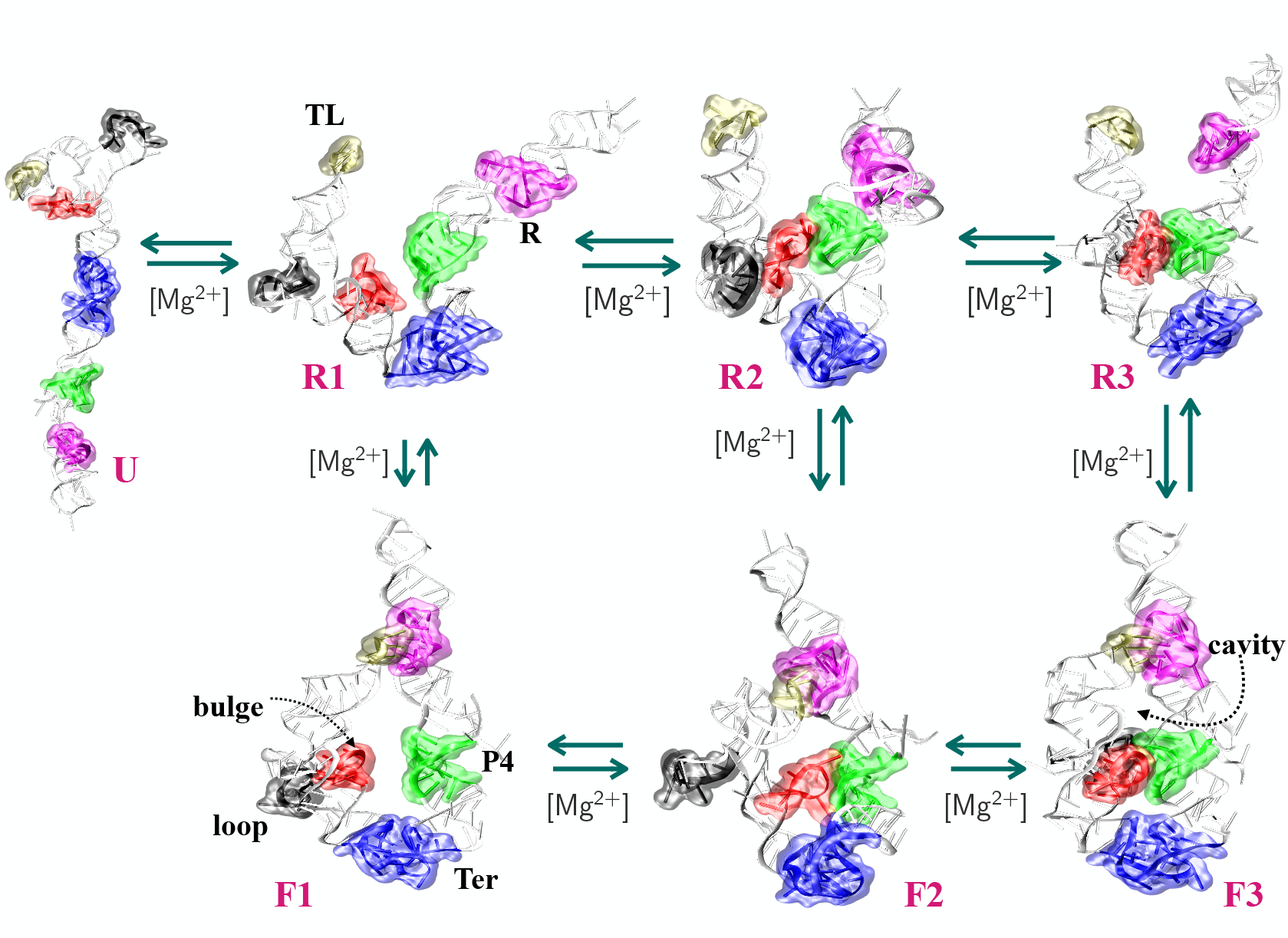
Representative conformations of seven states (U, R1, R2, R3, F1, F2, and F3) and their tertiary interactions are highlighted in different colors. The tertiary interactions TL and R are shown in the R1 structure. Bulge, loop, P4 and Ter interactions are shown in the F1 structure. Cavity is shown in the F3 structure. All the tertiary interactions are formed due to the condensation of Mg^2+^ ions.

**Figure 3:**
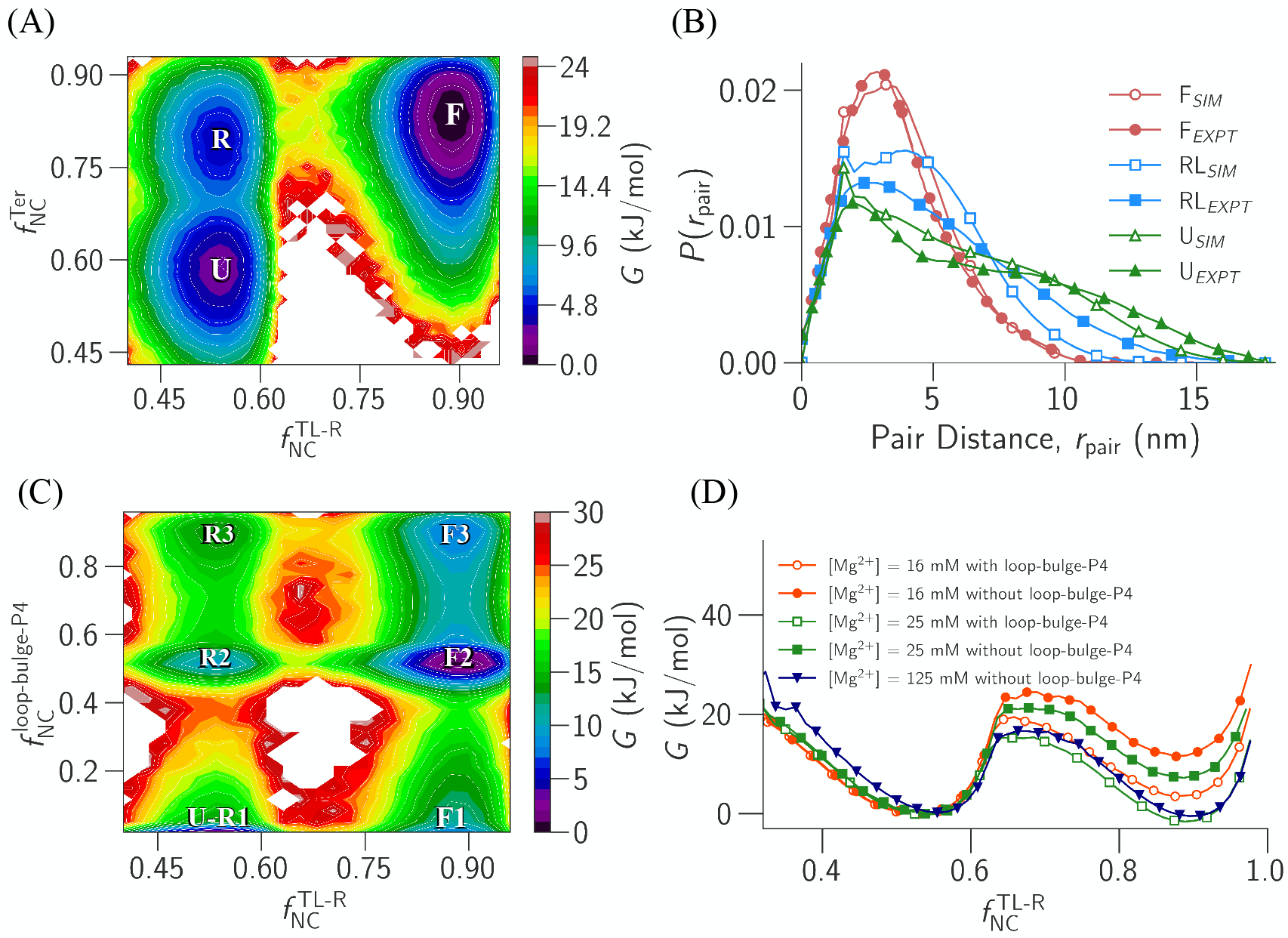
(A) The 2D FES of the intron projected on 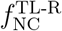 and 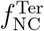 at [Mg^2+^] = 25 mM shows three distinct ensemble (U, RL and F). (B) Comparison of the probability distribution of phosphate–phosphate pair distances (*r*_pair_) for the three ensembles (U, RL, and F) obtained from CG simulations with experimental data (C) The 2D FES of the intron projected on 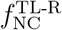 and 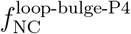 at [Mg^2+^] = 25 mM shows that the intron populates six states (U-R1, R2, R3, F1, F2 and F3) in its folding pathway. (D) FES projected on 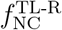 for different [Mg^2+^] with and without the presence of loop-bulge-P4 H-bonds.

The intron folding proceeds as follows. From the U state, the terminal interactions form first, giving rise to the R1 state (Fig. 2). From the R1 state, two routes are available: formation of the bulge-P4 contact leads to the R2 state, whereas the formation of the TL-R interaction leads to the F1 state. Now, F1 state can engage in bulge-P4 interaction, resulting in the transition to the F2 state. Subsequently, from the R2 state, the loop region can interact with the bulge-P4 region, forming the loop-bulge-P4 contact that characterizes the R3 state. Alternatively, the TL-R complex can form directly from R2, bypassing the loop-bulge-P4 interaction to form the F2 state. Eventually, both the F2 state (which forms loop-bulge-P4 interactions) and the R3 state (which forms TL-R docking) fold to the fully folded F3 state (Fig. 2).

The 2D free energy surfaces (FES) projected onto 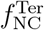 and 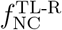 reveals three distinct states: (i) the U state 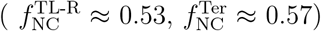, (ii) the RL state 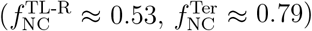 where only the terminal interactions are formed, and (iii) the F state 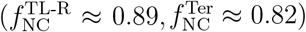 where the terminal interactions and the TL-R complex are formed (Fig. 3A, S5). The FES shows the U ↔ RL and RL ↔ F transitions. The probability distributions of phosphate–phosphate pair distances, *P* (*r*_pair_), for these three ensembles are in good qualitative agreement with the SAXS measurements of the P4–P6 domain (Fig. 3B).^59^

To resolve the substates of RL and F states, we projected the 2D FES onto 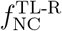 and 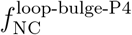 at [Mg^2+^] = 25 mM (Fig. 3C, 2, S3B). The FES reveals six well-defined states. The U-R1 region 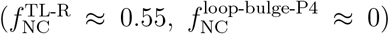 includes structures that may or may not contain terminal interactions but lack both loop-bulge-P4 and TL-R contacts. Thus, it contains R1 along with the U state. The R2 state 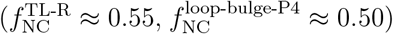 features the bulge-P4 interaction while remaining undocked at the TL-R interface. The R3 state 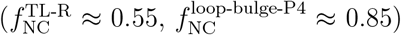 contains the fully formed loop-bulge-P4 contact but no TL-R docking. In the F1 state 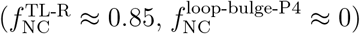, the TL-R complex is formed while the loop-bulge-P4 interaction is absent. In the F2 state 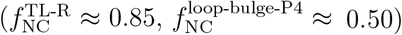, both the TL-R complex and bulge-P4 interaction are forming. Finally, the F3 state 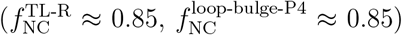 corresponds to the fully folded conformation, where all major tertiary contacts—including both TL-R docking and loop-bulge-P4 interactions—are established (Fig. 1B, 3A,B). Terminal interactions are present in R1, R2, R3, F1, F2, and F3. The contact maps for all these six states reveal the specific nucleotide–nucleotide interactions that characterize each state (Fig. S6).

The relative stabilities of the states at [Mg^2+^] = 25 mM follow the order - R3 < R2 ≈ F1 < F3 < U-R1 < F2 (Fig. 3C). We observed that the native-like F3 state is not the most stable, but the F2 state, which is close to the native conformation, is globally stable. However, we observed that as we increased [Mg^2+^] from 25 to 125 mM, the stability of the F3 state increased as the loop-bulge-P4 tertiary interaction 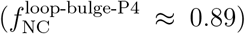 gained stability, but the F3 state was still not the global minimum (Fig. S7). The loop-P4 tertiary interactions are stabilized by three noncanonical H-bonds (two base-base and one phosphate-sugar H-bonds)(Fig. S8A,B). The additional stability is from several Mg^2+^ ions that are bound to the loop-bulge-P4 region through multiple inner-shell (IS) and outer-shell (OS) interactions contributing to the stability of this tertiary interaction (Fig. S9). However, these specific water-mediated interactions are absent in the CG TIS model, as water molecules are implicitly accounted for in this model. Hence, the F3 state is less stable than the F2 state in the CG simulations.

### Loop-Bulge-P4 Tertiary Interactions Stabilize the Docked TL-R Complex

To probe the importance of the loop-bulge-P4 tertiary interaction (C64-A70, U80-G86, U5-G9, and A108-A112) in facilitating the docking in TL-R complex, we performed CG simulations in which hydrogen bonding interactions within the loop-bulge-P4 region were disabled by turning off their interaction potentials (Table S3). Simulations were carried out at [Mg^2+^] = 16, 25, and 125 mM, while maintaining a constant background [Na^+^] = 150 mM. We projected the FES on 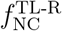 to check the stability of the docked state of the TL-R complex (Fig. 3D). The simulations show that the stability of the docked state of the TL-R complex decreased when the loop-bulge-P4 hydrogen bonds are disabled compared to when the hydrogen bonds are present. Interestingly, when we significantly increased the [Mg^2+^] (≈ 125 mM), we observed that both the undocked and the docked states are showing almost similar stability even when the loop-bulge-P4 hydrogen bonds are disabled (Fig. 3D). Therefore, we conclude that indeed, disruption of the loop-bulge-P4 interactions affects the intron folding at physiological [Mg^2+^]. However, at elevated [Mg^2+^] (≈ 125 mM), the loss of loop-bulge-P4 hydrogen bonds is compensated by Mg^2+^-mediated interactions, which restore the intron folding (Fig. 3D). This behavior is in agreement with the experimental observations by Murphy et. al.^27,92^

In the presence of loop-bulge-P4 tertiary interactions, the dominant intron folding pathways for [Mg^2+^] = 12−32 mM are: (i) U-R1 → R2 → R3 → F3 and (ii) U-R1 → R2 → F2 → F3 (Fig. 2). The least favored folding route is U-R1 → F1 → F2 → F3 (Fig. 2). Previous smFRET experiments^60,61^ also support the same preferred folding pathway of the intron, showing that from the elongated U state, the loop-bulge-P4 or bulge-P4 tertiary contacts form first, thereby facilitating the subsequent docking of the TL-R complex. These findings suggest that the loop-bulge-P4 or bulge-P4 interactions facilitate coaxial stacking between the P5a and P5b helices, which reduces the conformational entropy of TL and R helices, leading to the efficient formation of the TL-R docked complex (Fig. 1B).

### Mg^2+^ Ions Influence Tertiary Contact Formation in the Intron

We computed the local ion concentration of Mg^2+^ 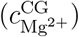 around the phosphate beads for the six states (U-R1, R2, R3, F1, F2, and F3) at [Mg^2+^] = 25 mM (see Data Analysis in SI) to understand the role of Mg^2+^ ions for the formation of RL and F ensembles in intron folding (Fig. 4, S10). The 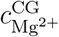 profiles show broadly similar peak distributions for all states, reflecting the formation of secondary structure (Fig. S10). However, there are differences at the TL-R complex and loop-bulge-P4 region, where tertiary interactions are involved (Fig. 4A,B) depending upon the states (Fig. 2, 3C).

**Figure 4:**
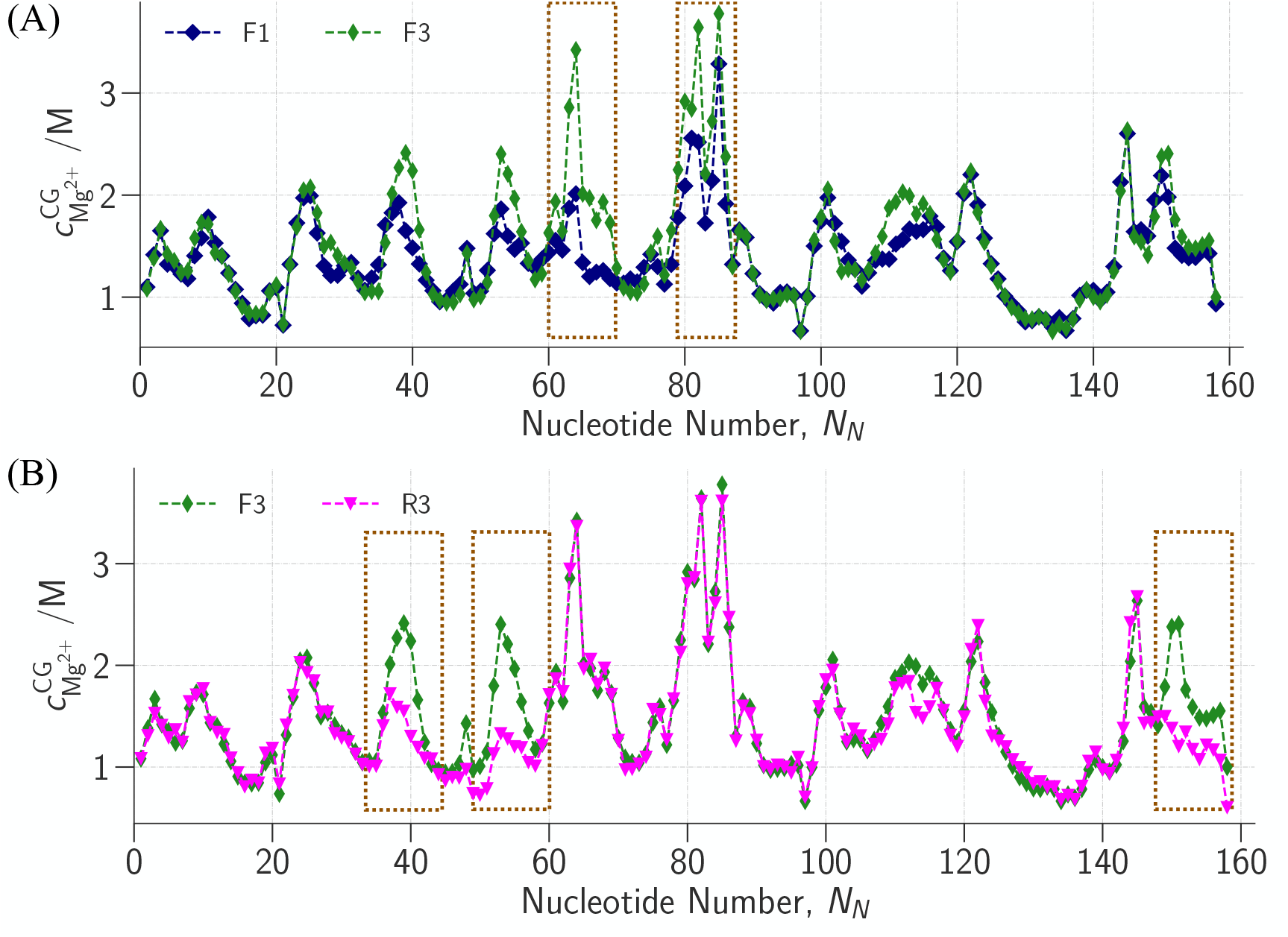
Local Mg^2+^ concentration 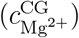 around the phosphate beads in (A) F1 and F3 states, (B) R3 and F3 states.

The comparison of 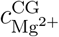 of state U-R1 with R2 and state F1 with F2, show that Mg^2+^ ions condensed more in the A82-A84 region for states R2 and F2, respectively, facilitating the formation of bulge-P4 interaction (Fig. S11A,B). Whereas, the comparison of 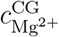 of state R2 with R3 and state F2 with F3 shows that Mg^2+^ ions condensed preferentially in the U60-A70 region for states R3 and F3, respectively, indicating formation of loop-P4 interactions (Fig. S11C, S12A). Hence, in the transition from F1→F3 and U-R1→R3, we observed Mg^2+^ ions are condensed more in the U60-A70 and A82-A84 promoting the formation of loop-bulge-P4 interactions (Fig. 4A, S12B). This indicates that in order to form the loop-bulge-P4 domain, Mg^2+^ ions need to be condensed in the loop-bulge-P4 domain. This is in agreement with the NMR, ^62^ smFRET,^60^ and chemical footprinting^93^ studies where they referred to the native loop-bulge-P4 domain as Mg^2+^ ion binding region.

The comparison of 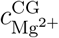 of state U-R1 with F1 and state R2 with F2 shows that Mg^2+^ ions mainly condensed in the cavity region beneath TL-R complex (G47-A57, G149-U157), facilitating the formation of TL-R docked complex (Fig. S13A,B and 2). In contrast, comparison between state R3 and F3, high Mg^2+^ ion condensation is observed not only at G47-A57 and G149-U157 but also around nucleotides A37-C41. This indicates that in addition to supporting the TL-R docking, Mg^2+^ ions also aid in forming a cavity beneath the TL-R complex during the R3→F3 transition (Fig. 4B, 2). Therefore, the formation of the loop-bulge-P4 interaction helps to stabilize the cavity region below the TL-R docked complex.

### TL-R Docking-Undocking Pathway Reveals Three Distinct States

The limitation of the CG TIS RNA model is that it cannot capture water-mediated interactions between Mg^2+^ ions and electronegative RNA heavy atoms due to the lack of explicit solvent water molecules. As a result, as shown in the previous sections, state F2 is more stable than state F3 in CG simulations due to the absence of water-mediated interactions that are critical for stabilizing the F3 state (Fig. 3C). FRET studies also have shown that Mg^2+^ ions stabilize the docked TL-R complex. ^40^

To investigate the R3 (undocked) ⇌ F3 (docked) transition and the role of hydrated Mg^2+^ ions on this transition (Fig. 2), we performed well-tempered metadynamics (WT-MetaD) simulations using all-atom RNA models.^79^ In WT-MetaD, we chose two CVs - (i) center of mass distance between the GAAA tetraloop (TL) and receptor (R), *d*_*com*_, and (ii) native contacts of TL-R complex (inter-helix) 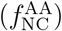 (Table S5) (See Methods and Data Analysis in the SI). In the WT-MetaD simulations, we observed multiple transitions between the docked and undocked conformations, indicating good sampling of the CV space (Fig. S14).

The 2D FES projected on *d*_*com*_ and 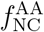 shows three distinct minima: (i) F3 state 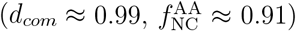, (ii) R3(1) state 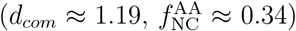, and (iii) R3(2) state 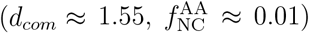 (Fig. 5A). The R3 state observed in the CG simulations (Fig. 2, 3C), where the TL-R docking is not observed, is split into two non-native intermediate states, R3(1) and R3(2), in the all-atom simulations (Fig. 5A). In the three states (F3, R3(1), and R3(2)), the interactions involving the TL-R complex differ, while the interactions between the bulge region and the P4 helix and tertiary interactions at the terminal region are stable (Fig. 5B, 2).

**Figure 5:**
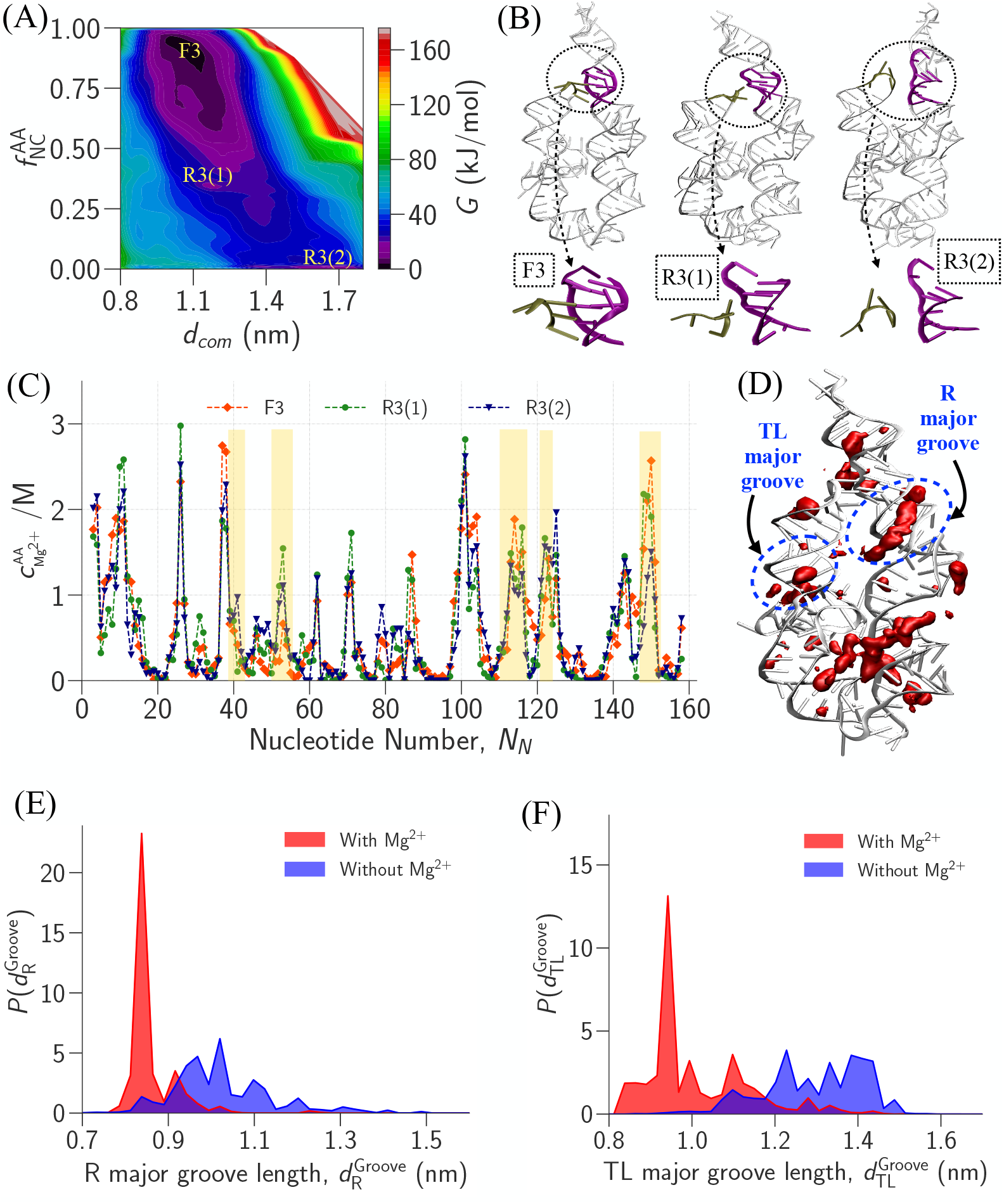
(A) The 2D FES projected on 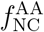 (inter native contacts between helices of GAAA tetraloop and receptor) and *d*_*com*_ (center of mass distance between two helices). The intron populates three states: F3, R3(1), and R3(2). (B) Representative snapshots are shown for three states where TL and R nucleotides are marked in olive and purple, respectively. (C) Local Mg^2+^ concentration 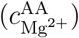 around the phosphate oxygens for F3, R3(1) and R3(2) states. Important regions are shaded in yellow. (D) Spatial density map of Mg^2+^ ions (red isosurfaces) around the phosphate backbone for F3 state. The isosurface value (ISO = 0.028) represents the fraction of time Mg^2+^ ions occupy the region enclosed by the surface. Probability distribution of major groove length of (E) R helix 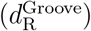 and (F) TL helix 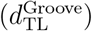 in the presence (red) and absence (blue) of Mg^2+^ condensation.

In the F3 state, where the TL and R helices are docked, in addition to the triplex H−bonding interaction between the nucleobases A49, U122, and A146, there are two additional hydrogen bonds between A123 and A124 (Fig. S15A). In contrast, in the R3(1) and R3(2) states−where the TL and R helices are undocked−instead of the triplex H-bonds, we observe a new H-bond between U147 and U122, and H-bonds between A123−A124 are broken (Fig. S15B). This suggests that the hydrogen-bonding network within the R helix is modulated by its docking status with the TL helix. This observation is consistent with the experimental findings, ^94^ further validating the results from the WT-MetaD simulations.

### Water-Mediated Mg^2+^-RNA OS Interactions Stabilize Non-Native Intermediates

As discussed in the previous section, although hydrogen-bonding networks between the TL and R helices are essential for TL-R docking (Fig. S15A,B), these interactions are absent in the intermediate states R3(1) and R3(2). This prompts the question: how do these intermediates maintain stability without such interactions? To address this, we examined whether hydrated Mg^2+^ ions contribute to their stabilization. We computed the local concentration 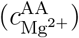 and spatial distribution of Mg^2+^ ions around the phosphate oxygen atoms in different states to understand the effect of Mg^2+^ ions on the docking-undocking transition of the TL-R complex in the WT-MetaD simulations (see Data Analysis in SI) (Fig. 5C,D and S16).

In the R3(1) state, 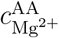 is high near the phosphate oxygen atoms of nucleotides A52, U53, U122, and A150, located at the TL-R complex due to the outer-shell (OS) coordination between these nucleotides and Mg^2+^ ions (Fig. 5C and S17A,B). Although native H-bonds are disrupted in the R3(1) state, the TL and R helices are in proximity due to these bridging OS interactions (Fig. 5B). In the R3(2) state, the TL helix moves further away from the R helix compared to the R3(1) state (Fig. 5B). In the R3(2) state, 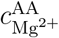 has peaks at U40, C41, C52 and U53 (Fig. 5C), and the interaction between TL and R helix is stabilized by OS bridging interactions between these nucleotides and the Mg^2+^ ions (Fig. S18). Although intermediate states are stabilized by the OS interactions between Mg^2+^ ions and RNA-phosphate-oxygens, the F3 state is more stable than the R3(1) and R3(2) states because of the formation of native like H-bonds in the TL-R region (Fig. S19A,B). Spatial distribution analysis of Mg^2+^ ions in the F3, R3(1), and R3(2) states show that Mg^2+^ ions condense in the major grooves of both the TL and R helices in all the states (Fig. 5D and S16). Similarly, Mg^2+^ ions also condensed in the major grooves of the P4 and P5a helices in all the states. These observations are in agreement with a previous simulation study. ^95^

We observed TL-R docked structures in the F3 basin showing native-like hydrogen-bonds, which are absent in the R3(1) and R3(2) states (Fig. S19A,B). In the broad F3 basin, we found intron conformations with the triplex H-bonding and conformations without the triplex H-bonding. In the conformations with the triplex H-bonding (A49-U122-A146), additional inter-helix H-bonds, A50-G148, A50-U122, and A51-C121, are observed between the nucleotides of TL and R helices (Fig. S19A). In contrast, in the absence of triplex H-bonding, the inter-helix hydrogen bonds are observed between A50-U122 and A51-G148 (Fig. S19B). Although in the F3 basin, the docked structures do not form the complete set of native hydrogen bonds reported in the crystal structure, the broken H-bonds are stabilized through water-mediated outer-shell (OS) interactions involving Mg^2+^ ions and the phosphate oxygens of A50, A51 and A123 of the TL and R helices (Fig. S20A). These Mg^2+^ ions are found in the spatial distribution of Mg^2+^ ions and 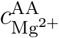 in the F3 basin (Fig. 5C,D).

In the F3 state, prominent peaks in 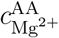 at nucleotides A112-A116, G149-U151 - corresponding to the major groove of the R helix - are significantly higher compared to those in the R3(1) and R3(2) states (Fig. 5C). The spatial distribution of Mg^2+^ ions also confirmed greater condensation of Mg^2+^ ions in the F3 state in this region compared to the intermediates (R3(1) and R3(2) states), which are stabilized by outer-shell (OS) interactions with these nucleotides (Fig. S20B). This suggests that Mg^2+^ ion condensation in the major groove is essential for stabilizing the docked state.

### Mg^2+^ Condensation Controls the Fluctuation in the Groove Length in Helices Critical for TL-R Docking

To test the hypothesis that condensation of Mg^2+^ ions in a helical groove is essential for the helix to maintain a critical length necessary for docking, we measured the major groove lengths by calculating the distances between selected phosphate atoms: (i) distance between phosphates C114 and A150 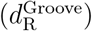 for the R helix, and (ii) distance between phosphates A37 and A57 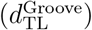 for the TL helix. We computed the probability distributions of these distances under two conditions in the R3(2) state: (i) when Mg^2+^ ions are condensed in the major grooves of R and TL helices, and (ii) when Mg^2+^ ion condensation is absent in the major grooves of R and TL helices (Fig. 5E,F).

The presence of Mg^2+^ ion condensation in the major groove of R helix results in a sharper distribution of 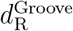 with an average length of 0.87 nm, while in the absence of Mg^2+^ ion condensation, the distribution is broad with an average of 1.04 nm (Fig. 5E). Similarly, for the TL helix, the distribution of 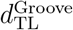 is sharp with an average 1.13 nm when Mg^2+^ ion condensation is observed and the distribution is broad with an average 1.26 nm in the absence of Mg^2+^ ion condensation (Fig. 5F). These results indicate that condensation of Mg^2+^ ions in the helical grooves stiffen the helices and reduce the fluctuations in their length, which will facilitate their docking. CG simulations further support this, showing similar Mg^2+^ ion condensation in the major grooves of the TL and R helices in the folded F3 state (Fig. S21). Comparable Mg^2+^ ion-mediated stabilization has been observed in DNA systems, where Mg^2+^ ions reduce fluctuations in the phosphate backbone and influence hydration patterns within the major groove. ^96,97^

Based on the spatial ion distribution and 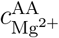 values, Mg^2+^ ion condensation is greater in the R helix in the F3 states compared to the R3(2) state (Fig. 5C,D). This enhanced Mg^2+^ ion condensation is likely driven by OS interactions between Mg^2+^ ions and nucleotides A112, C114, A116, U151, and G149 (Fig. S20B). These findings suggest that it is more critical for the R helix to be stiff and maintain a proper length^98^ compared to the TL helix for the correct docking and the formation of TL-R complex (F3 state).

## Conclusion

Using the P4-P6 domain in the *Tetrahymena thermophila* group I intron as a model system, this study illustrates the mechanism of Mg^2+^ ion-mediated tertiary interaction formation between a tetraloop and receptor, which is a prime tertiary interaction that enables the formation of long-range contacts in folded RNA structures. The simulations have revealed that the intron folds via seven distinct states (U, R1, R2, R3, F1, F2, and F3) with varying tertiary contacts. Among these states, several pivotal intermediates (e.g., R2, R3, F1, F2) strongly bias the selection of the folding pathway. We further showed that [Mg^2+^] higher than physiological limits can nullify the effect of loop-bulge-P4 interaction for the TL-R docking. This suggests that the strategic arrangement of conserved sequences participating in the loop-bulge-P4 interaction is essential to promote TL-R docking at physiological Mg^2+^ ion levels.^27,69,92^ Moreover, we identified that Mg^2+^ ions form OS contacts with the RNA phosphate oxygen of the TL and R helices to stabilize several non-native intermediate states across the folding pathway during the TL-R docking-undocking process. Interestingly, we also find that the condensation of Mg^2+^ ions in the grooves of helices containing TL and R is crucial for maintaining their stiffness and facilitating efficient docking. Without Mg^2+^ ion condensation, the helices become flexible, and the docking is inefficient. This site-specific Mg^2+^ ion condensation links local groove stabilization to the global tertiary architecture of RNA. Overall, these findings show that in larger RNA systems, proximal RNA-RNA interactions can play a pivotal role in maneuvering the global structural motion, rendering an optimal pathway for distal RNA-RNA interactions in physiological [Mg^2+^], and these insights into the RNA folding mechanisms have implications for ribozyme engineering. Several interesting questions remain in understanding the function of group I introns such as how do these intermediates contribute to their catalytic activity? Is it by facilitating structural transitions or stabilizing key conformations during folding?

## Supporting Information

Data analysis; CG parameters (Table S1, S2, S3, and S4); Different native contacts (Table S5); Docked and undocked TL-R complex (Fig. S1); Convergence for WT-MetaD simulations (Fig. S2); Change of CVs with time for CG simulation (Fig. S3); FES of different CVs (Fig. S4); Snapshots of U, RL and F states (Fig. S5); Contact map for six states (Fig. S6); FES projected onto 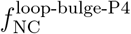 (Fig. S7); Crystal H-bonds in loop-P4 region (Fig. S8); Native IS and OS interactions (Fig. S9); Local ion concentrations (Fig. S10−S13); Change of CVs with time for WT-MetaD simulation (Fig. S14); H-bonding arrangement during docking and undocking (Fig. S15); Native like H-bonds in docked TL-R complex (Fig. S19); Spatial density maps for different states (Fig. S16 and S21); OS interactions in R3(1), R3(2) and F3 states (Fig. S17, S18, and S20).

## Acknowledgement

GR acknowledges funding from the Science and Engineering Research Board (SERB) through the grant CRG/2023/002817. SH acknowledges the Prime Minister’s Research Fellowship (PMRF). DM acknowledges the research fellowship from the Indian Institute of Science, Bangalore. We acknowledge the National Supercomputing Mission (NSM) for providing computing resources of “Param Pravega” at IISc and “Param Brahma” at IISER Pune, supported by the Ministry of Electronics and Information Technology (MeitY) and the Department of Science and Technology (DST), Government of India.

