## Supplementary figures and tables for "Mg^2+^-Dependent Multistep Folding and Stabilization of the GAAA Tetraloop-Receptor Interaction in a Group I Intron"

### Data Analysis

#### Fraction of Native Contacts

The native contacts in the intron are computed using the TIS model of the folded crystal structure. A pair of beads $i$ and $j$ have a native contact when their sequence separation ($|i-j|$) exceeds 10 beads and the distance between them ($r_{ij}$) is less than 15 Å in the crystal structure. The fraction of native contacts ($f_{\mathrm{NC}}$) for the $k^{th}$ conformation is calculated using the equation

$$f_{\mathrm{NC}}^{(k)} = \frac{N_{\mathrm{NC}}^{(k)}}{N_{\mathrm{NC}}^{(\mathrm{cry})}} \tag{S1}$$

where $N_{\mathrm{NC}}^{(k)}$ and $N_{\mathrm{NC}}^{(\mathrm{cry})}$ are the number of native contacts present in the $k^{th}$ conformation and crystal structure, respectively.

#### Free Energy Surface (FES) from CG Simulations

We projected the free energy surface (FES) onto $f_{\mathrm{NC}}$ to identify the various thermodynamic states populated during the intron folding. We computed the FES, $G(f_{\mathrm{NC}})$ using the equation

$$G(f_{\mathrm{NC}}) = -k_{\mathrm{B}}T \ln\big(P(f_{\mathrm{NC}})\big) \tag{S2}$$

where $k_{\mathrm{B}}$ is the Boltzmann constant, and $P(f_{\mathrm{NC}})$ $\big(= \int \mathrm{d}\boldsymbol{r}\ \delta[f_{\mathrm{NC}} - f_{\mathrm{NC}}(\boldsymbol{r})]P(\boldsymbol{r})\big)$ is the probability that RNA conformations given by coordinates $\boldsymbol{r}$ have the fraction of native contacts value $f_{\mathrm{NC}}$.

We calculated the fraction of native contacts in the full P4-P6 domain ($f_{\mathrm{NC}}$), within the TL-R complex ($f_{\mathrm{NC}}^{\mathrm{TL\text{-}R}}$), within the terminal region ($f_{\mathrm{NC}}^{\mathrm{Ter}}$), and between the L5c loop, the major bulge, and the P4 helix ($f_{\mathrm{NC}}^{\mathrm{loop\text{-}bulge\text{-}P4}}$) in the CG simulations (Table S5).

#### Radius of Gyration

The radius of gyration ($R_g$) of the intron conformations was calculated using the equation

$$R_{\mathrm{g}} = \left( \frac{\sum_{i=1}^{N} m_i \vec{r}_i^{\,2}}{\sum_{i=1}^{N} m_i} \right)^{1/2} \tag{S3}$$

where $m_i$ is the mass of the $i^{\mathrm{th}}$ coarse-grained bead, and $\vec{r}_i$ is the distance of the $i^{\mathrm{th}}$ bead from the center of mass of the conformation. The mass values of all the bead types are listed in Table S4.

#### Local Ion Concentration Around RNA

To analyze ion condensation around RNA, we computed the local concentration ($c_j$) of divalent metal ions near the phosphate oxygen atoms (O1P and O2P) in all-atom (AA) simulations and phosphate beads in CG simulations. The local ion concentration was calculated using[1]

$$c_j = \frac{1}{N_{\mathrm{A}} V_c} \int_0^{r_c} \mathrm{d}r \; 4\pi r^2 \; \rho_j(r) \tag{S4}$$

where $\rho_j(r)$ is the number density of ion type $j$ at a distance $r$ from a given RNA atom, $V_c$ is the spherical volume within a cutoff radius $r_c$, and $N_{\mathrm{A}}$ is Avogadro number.

In AA simulations, the cutoff distance $r_c^{(\mathrm{AA})}$ is the sum of the ionic radius ($R_{\mathrm{M}^{2+}}$), the radius of RNA phosphate oxygen atoms (O1P and O2P), and the characteristic inner-shell contact distance ($R_{\mathrm{IS}}$). The values $R_{\mathrm{Mg}^{2+}} = 1.353$ Å, $R_{\mathrm{Co}^{2+}} = 1.288$ Å, $R_{\mathrm{O}} = R_{\mathrm{N}} = 1.7$ Å, and $R_{\mathrm{IS}} \approx 2.0$ Å, leading to $r_c^{(\mathrm{AA})} \approx 5$ Å.

In CG simulations, the cutoff distance is given by $r_c^{(\mathrm{CG})} = (R_{\mathrm{M}^{2+}} + R_{\mathrm{P}} + \Delta r)$, where $R_{\mathrm{P}} = 2.1$ Å is the radius of the CG phosphate bead and $\Delta r = 1.6$ Å is a margin distance to account for spatial resolution, resulting in $r_c^{(\mathrm{CG})} \approx 5$ Å. To maintain consistency between AA and CG systems, we uniformly used a cutoff radius of $r_c = 5$ Å for all local concentration calculations.

#### Spatial Distribution of Ions Around RNA

To visualize the distribution of $Mg^{2+}$ ions around RNA, we generated spatial density maps using the volmap plugin in VMD software.[2] All RNA conformations were structurally aligned to the final frame of the CG simulation trajectory to ensure consistent reference geometry. This alignment was carried out using the Kabsch algorithm,[3] as implemented in VMD, and the corresponding ion coordinates were transformed accordingly. Each metal ion was treated as a sphere with its van der Waals radius. A volumetric grid with a resolution of 10 $Å^3$ was defined around the RNA. For each conformation, grid points occupied by an $Mg^{2+}$ ion were assigned a value of 1, and the unoccupied grid points were assigned a value of 0. These binary values were then averaged over all frames to generate the final spatial ion distribution. We also used VMD for both structural superposition[4] and image rendering.[5]

Table S1: Watson–Crick and non-Watson–Crick base–base hydrogen bond interactions used in the CG model. The number of hydrogen bonds are given in the brackets.

| **Base–Base Hydrogen Bonds (Number of Hydrogen Bonds)** |
| --- |
| G47–C52 (3), G56–C43 (3), G58–C41 (3), G73–C63 (3), G72–C64 (3), G78–C36 (3), G79–C35 (3), G32–C87 (3), G89–C30 (3), G27–C91 (3), G92–C26 (3), G93–C25 (3), G98–C19 (3), G16–C101 (3), G15–C102 (3), G10–C106 (3), G9–C107 (3), G8–C109 (3), G110–C7 (3), G6–C111 (3), G149–C120 (3), G148–C121 (3), G143–C127 (3), G140–C130 (3), G132–C138 (3), G1–C115 (3), A44–U55 (2), A57–U42 (2), A59–U40 (2), A71–U65 (1), A34–U80 (2), A31–U88 (2), A90–U29 (2), A112–U5 (2), A3–U156 (2), A150–U119 (2), A124–U147 (2), A146–U122 (2), A144–U126 (2), A128–U142 (2), A129–U141 (2), A131–U139 (2), A133–U137 (2), A2–C114 (1), A3–G113 (2), A11–A105 (1), A12–A104 (1), A37–G62 (2), A38–G61 (2), A49–A146 (2), A51–G148 (1), A116–A117 (1), A123–A124 (1), C22–C95 (1), G1–A154 (1), G24–A94 (2), G48–A51 (1), G62–A76 (1), G79–A84 (1), U5–U157 (1), U33–A85 (2), U65–A71 (1), U66–G86 (2), U122–A146 (2), G14–U103 (1), G17–U100 (2), U18–G99 (2), G39–U60 (2), G45–U54 (2), G46–U53 (2), G118–U151 (2), G125–U145 (2), G99–C22 (1), G98–C95 (1), A2–G155 (1), A3–U156 (2), A82–G110 (1), A21–A96 (1) |

Table S2: Base-sugar, sugar-sugar, phosphate-sugar hydrogen bonds and base-base stacking interactions. Number of hydrogen bonds are given in brackets.

| **Base-Sugar** | **Sugar-Sugar** | **Phosphate-Sugar** | **Base-Base Stacking** |
|---|---|---|---|
| A11−A104 (1)<br>A20−A96 (1)<br>A50−G148 (1)<br>C111−A82 (1)<br>U151−A116 (1)<br>U145−A123 (1)<br>U66−G9 (2)<br>A84−G79 (2)<br>A84−G62 (2)<br>G24−G99 (1)<br>A50−G48 (1)<br>U80−A84 (1)<br>G152−U151 (1)<br>G62−C36 (1)<br>C7−A82 (1)<br>A82−G110 (1)<br>A81−G8 (1)<br>U83−A37 (1)<br>C35−A84 (1) | A3−U157 (1)<br>A37−U83 (1)<br>A50−U122 (1)<br>A51−C121 (1)<br>A51−U122 (1)<br>C7−A82 (1)<br>C35−A84 (1)<br>C52−G149 (1)<br>C95−G98 (1)<br>G24−U100 (1)<br>G8−A81 (1)<br>U80−G86 (1)<br>C95−G99 (1) | G110−G67 (1) | A12−A105 (1)<br>A34−A85 (1)<br>A49−A124 (1)<br>A76−G78 (1)<br>A96−G98 (1)<br>U65−A69 (1)<br>U66−A81 (1)<br>A11−A104 (1)<br>A20−A96 (1)<br>A116−U151 (1)<br>A123−U145 (1)<br>A123−A146 (1)<br>A146−G148 (1)<br>A84−A37 (1)<br>C22−G24 (1)<br>U80−A85 (1) |

Table S3: Base-base, base-sugar, sugar-sugar, and base-base stacking interactions present in loop-bulge-P4 tertiary interaction. Number of hydrogen bonds are given in brackets.

| **Base-Base** | **Base-Sugar** | **Sugar-Sugar** | **Base-Base Stacking** |
|---|---|---|---|
| A34−U80 (2)<br>G79−A84 (1)<br>U33−A85 (2)<br>U66−G86 (2)<br>A82−G110 (1) | C111−A82 (1)<br>U66−G9 (2)<br>A84−G79 (2)<br>A84−G62 (2)<br>U80−A84 (1) | A37−U83 (1)<br>C7−A82 (1)<br>C35−A84 (1)<br>G8−A81 (1)<br>U80−G86 (1) | A34−A85 (1)<br>U66−A81 (1)<br>A84−A37 (1)<br>U80−A85 (1) |

Table S4: RNA[1] and ions[6,7] parameters used in the CG simulations.

| Type | $R_i$ (Å) | $m_i$ (amu) | $\epsilon_i$ (kcal mol$^{-1}$) | $Z_i$ |
|---|---|---|---|---|
| P | 2.1 | 62.971 | 0.2 | -1 |
| S | 2.9 | 131.108 | 0.2 | 0 |
| A | 2.8 | 134.119 | 0.2 | 0 |
| G | 3.0 | 150.118 | 0.2 | 0 |
| C | 2.7 | 110.094 | 0.2 | 0 |
| U | 2.7 | 111.079 | 0.2 | 0 |
| $Mg^{2+}$ | 1.353 | 24.305 | 0.009 | +2 |
| $Na^{+}$ | 1.226 | 22.990 | 0.168 | +1 |
| $Cl^{-}$ | 2.760 | 35.453 | 0.012 | -1 |

Table S5: Number of native contacts in different tertiary structures of the P4-P6 domain of the intron

| **Native Contact** | **Number of Contacts** | **Associated Nucleotides** |
|---|---|---|
| $f_{\text{NC}}$ | 7037 | All nucleotides in P4-P6 domain |
| $f_{\text{NC}}^{\text{TL-R}}$ | 1027 | G48-A51, C120-G125, U145-G149 |
| $f_{\text{NC}}^{\text{loop-bulge-P4}}$ | 211 | G6-G8, U66-C68, A81-U83, C109-C111 |
| $f_{\text{NC}}^{\text{Ter}}$ | 1067 | G16-C25, G93-C101 |
| $f_{\text{NC}}^{\text{AA}}$ | 278 | Inter contacts between (G48-A51) and (C120-G125, U145-G149) |

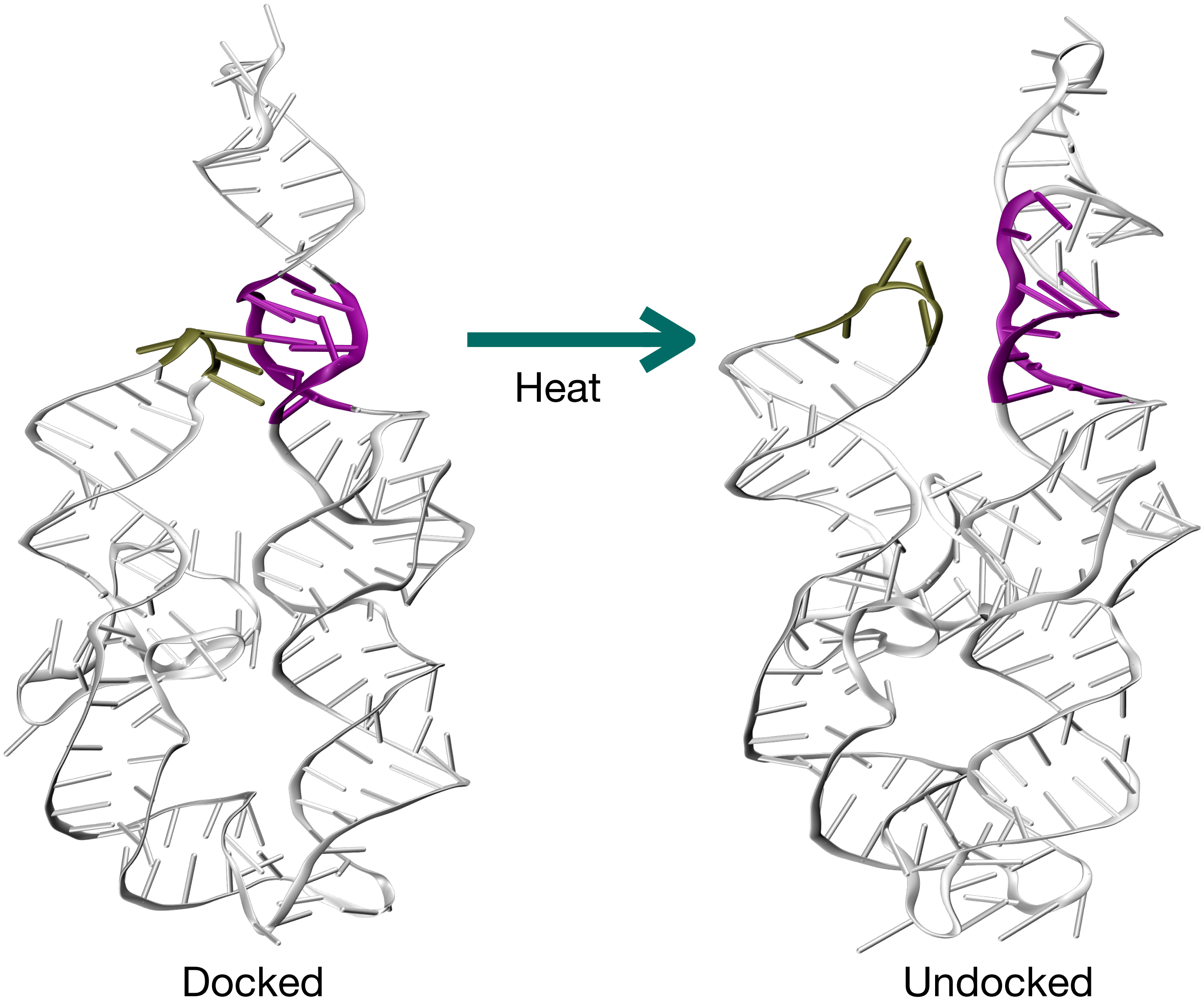


Figure S1: Representative structure of the undocked conformations obtained after heating the docked structure from 300 K to 350 K. TL (4 nt) and R (11 nt) regions are shown in olive and purple, respectively.

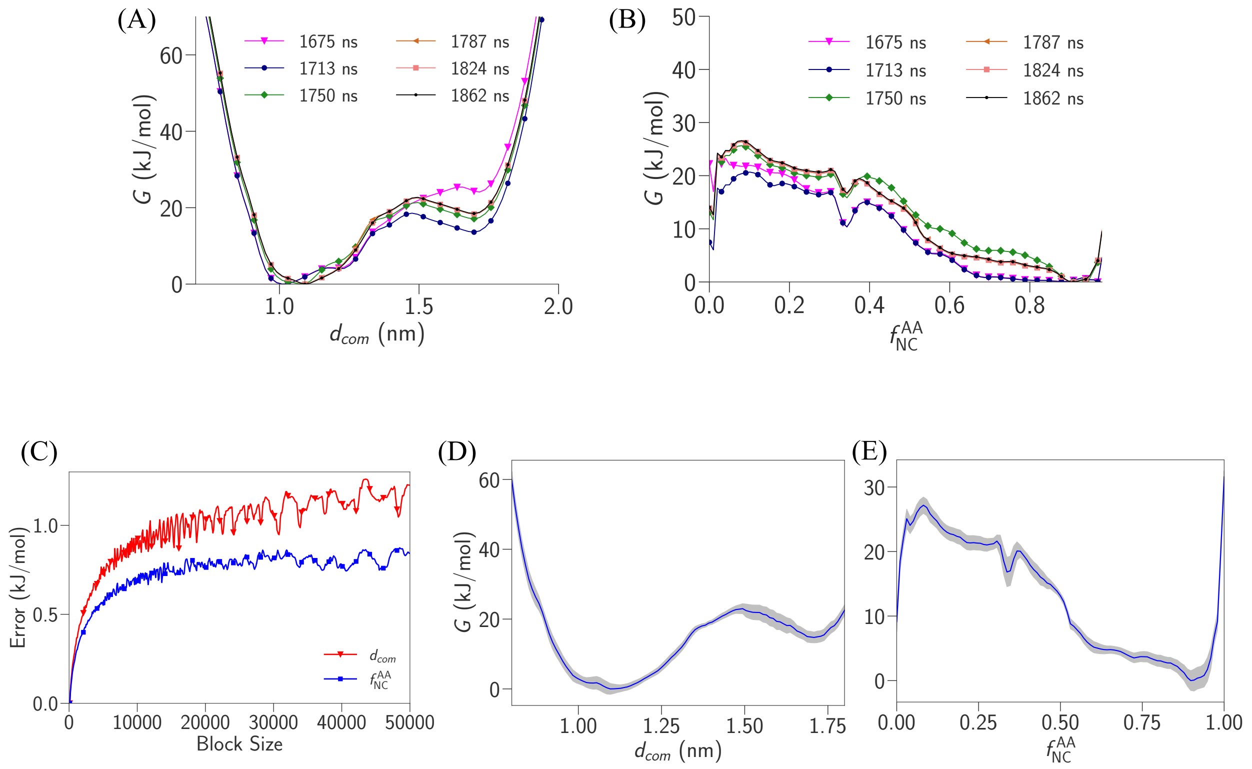


Figure S2: Convergence of WT-MetaD simulations: The free energy G as a function of (A) $d_{com}$ and (B) $f_{\mathrm{NC}}^{\mathrm{AA}}$. (C) Error bars for the CVs, $d_{com}$ and $f_{\mathrm{NC}}^{\mathrm{AA}}$ are computed using block analysis. The silver-shaded region represents the errors in the free energy profiles, and the blue curve indicates the average free energy profiles for (D) $d_{com}$ and (E) $f_{\mathrm{NC}}^{\mathrm{AA}}$.

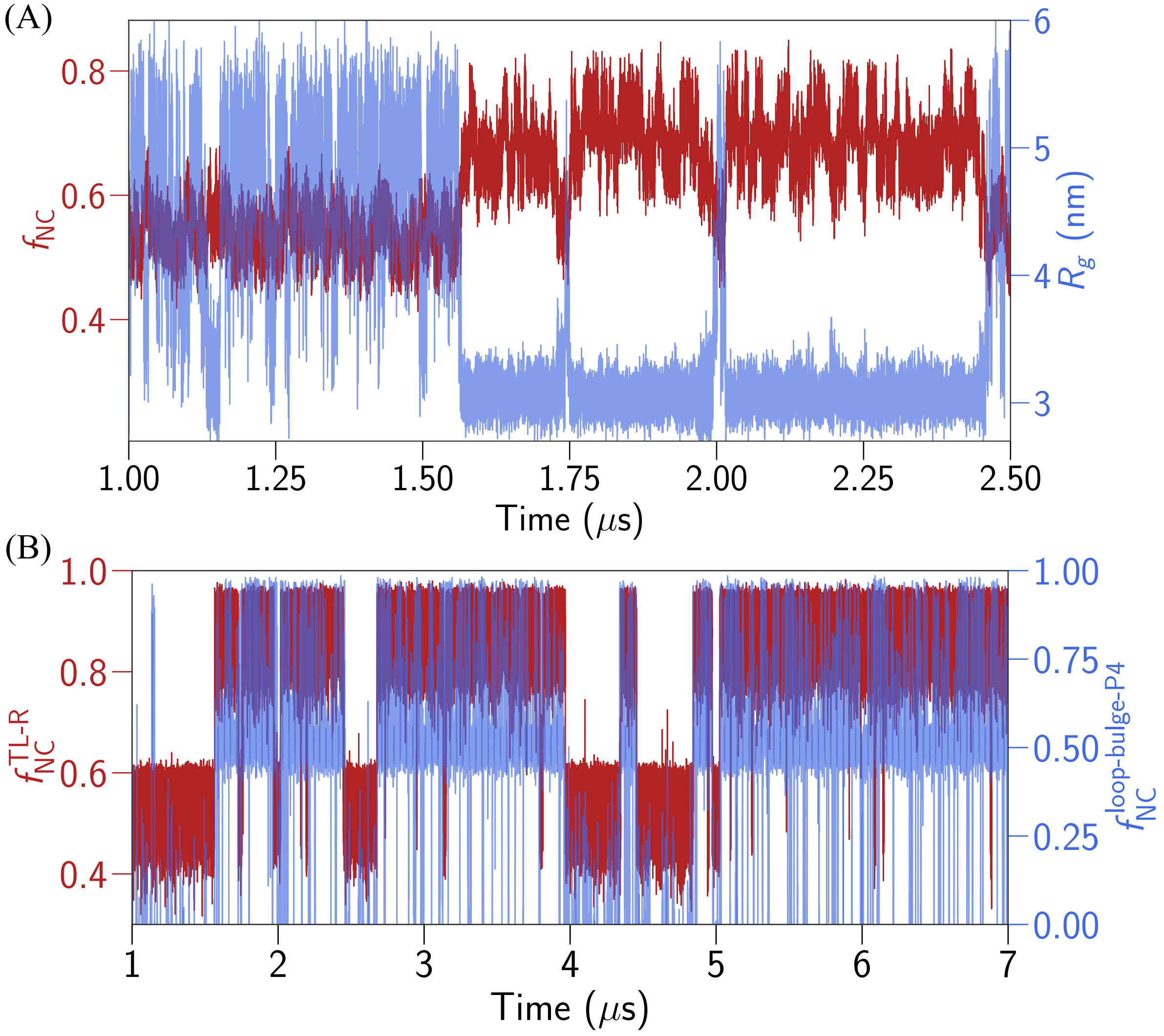


Figure S3: (A) $f_{\mathrm{NC}}$ and $R_g$ are plotted as a function time in a representative CG simulation for $[\mathrm{Mg}^{2+}] = 25$ mM. (B) $f_{\mathrm{NC}}^{\mathrm{TL\text{-}R}}$ and $f_{\mathrm{NC}}^{\mathrm{Bulge\text{-}P4}}$ are plotted as a function time in a representative CG simulation for $[\mathrm{Mg}^{2+}] = 25$ mM.

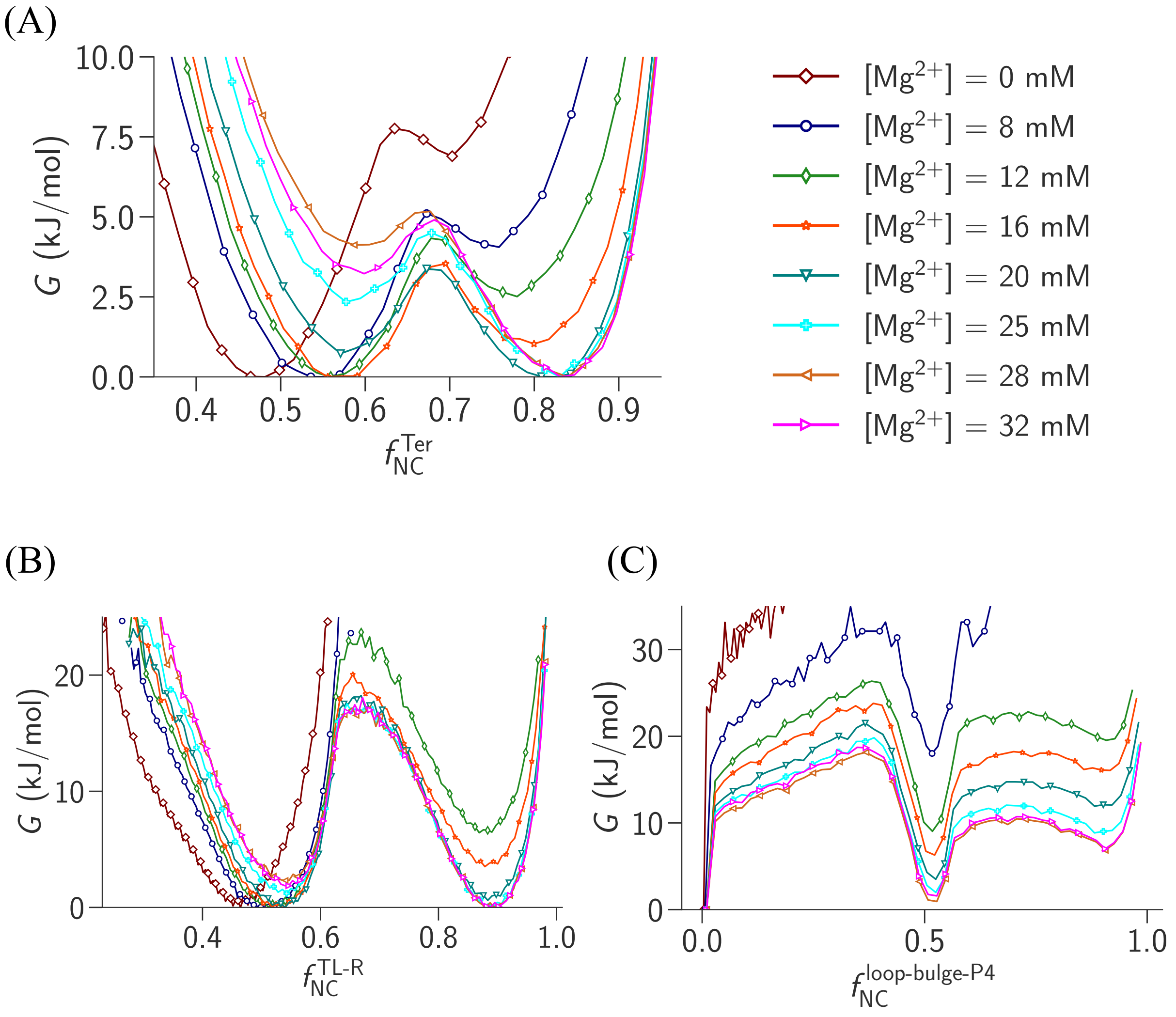


Figure S4: Free energy surface projected onto (A) $f_{NC}^{Ter}$, (B) $f_{NC}^{TL\text{-}R}$, and (C) $f_{NC}^{loop\text{-}bulge\text{-}P4}$ in CG simulations for different $[Mg^{2+}]$.

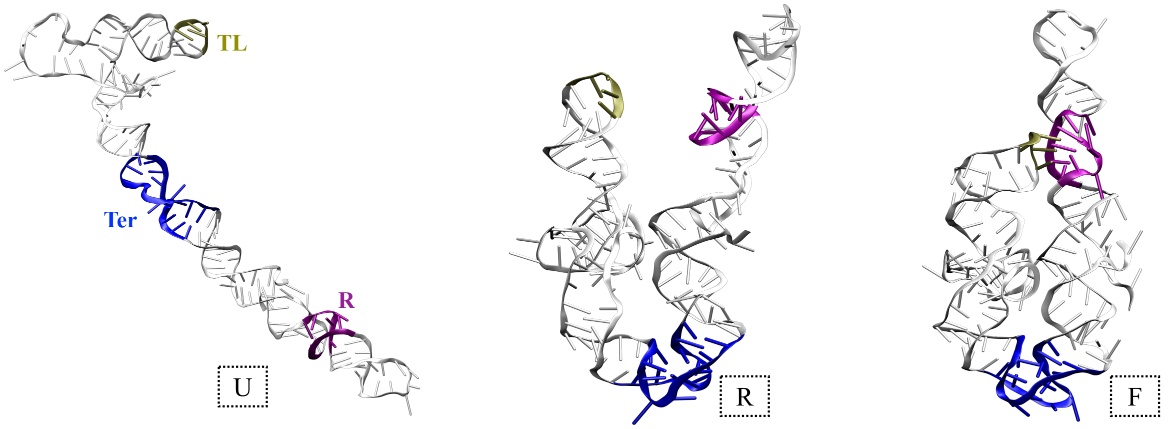


Figure S5: Representative snapshots of three states from representative CG simulation for $[Mg^{2+}] = 25$ mM: unfolded (U), relaxed (RL) and folded (F).

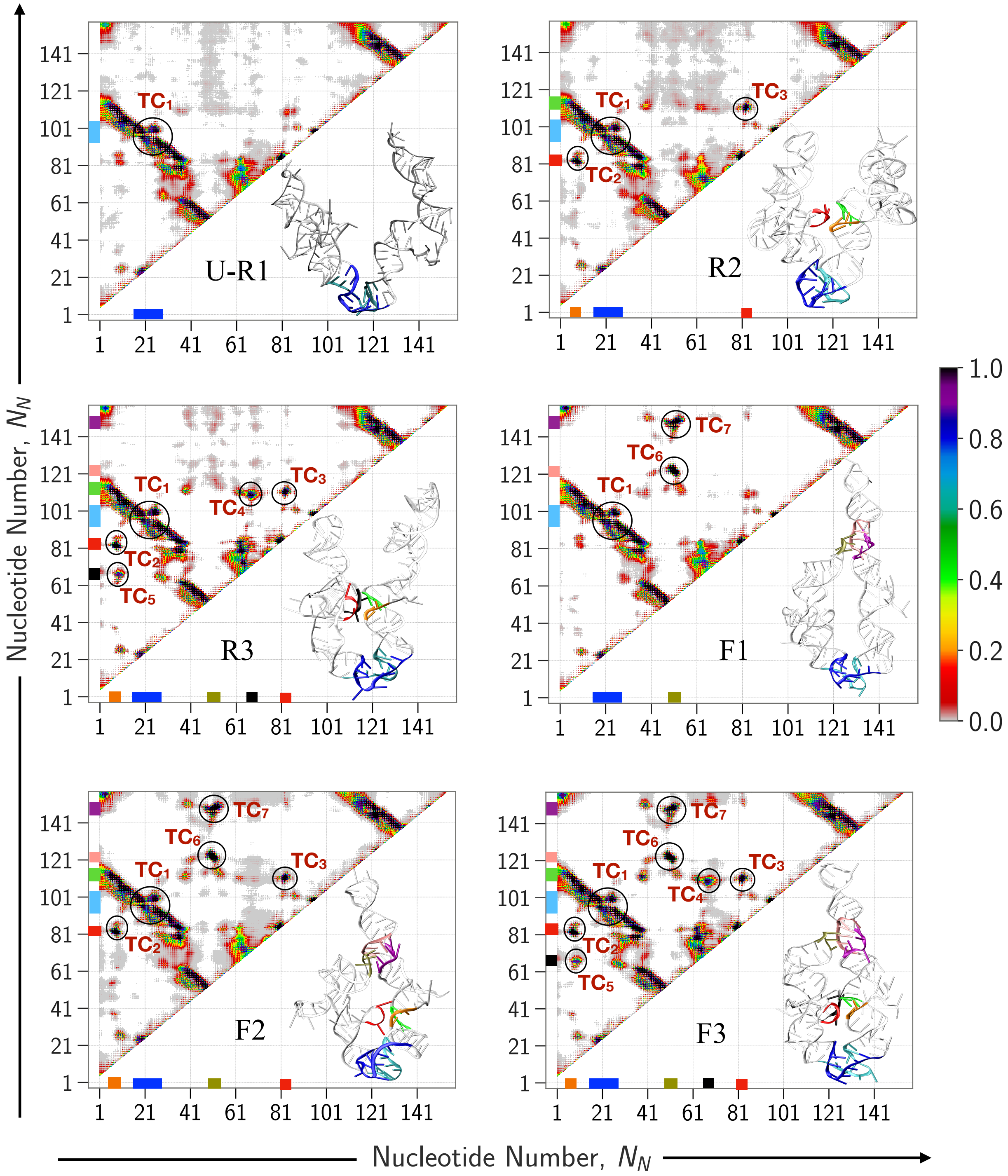


Figure S6: Average contact maps of the conformations in the states U-R1, R2, R3, F1, F2 and F3. Representative structures are shown for each state. The TL-R complex, loop-bulge-P4 region and and terminal parts on the representative structures are shown as follows: (1) TL-R Complex: G48-A51 (tan), C120-G125 (pink), U145-G149 (purple), (2) loop-bulge-P4 region: A81-U83 (red), C6-G8 (orange), U66-C68 (black), A109-A112 (green), and (3) terminal part: G16-C25 (blue), G93-C101 (cyan). The same color blocks on the $x$ and $y$ axes are a guide to the eye. The tertiary contacts, $TC_1$ to $TC_7$, are highlighted using black circles.

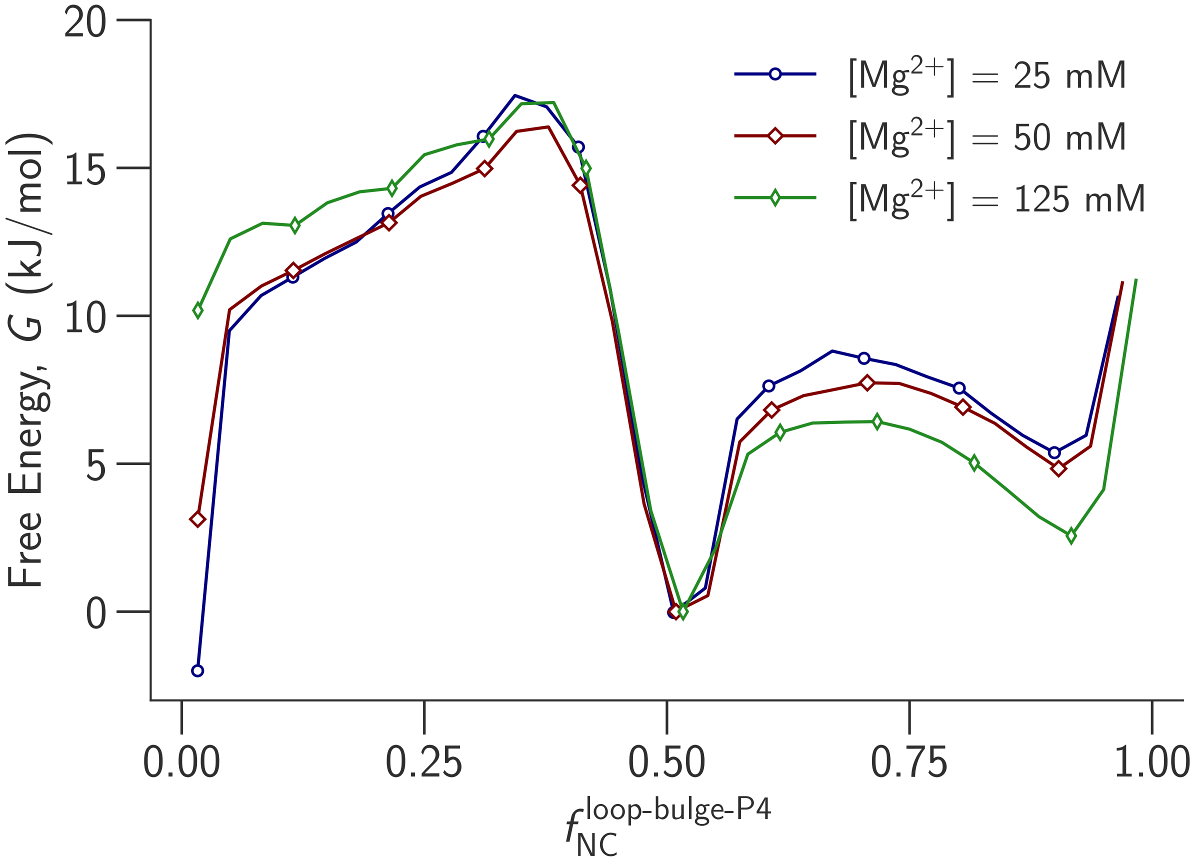


Figure S7: FES projected onto $f_{\mathrm{NC}}^{\text{loop-bulge-P4}}$ for different [$Mg^{2+}$].

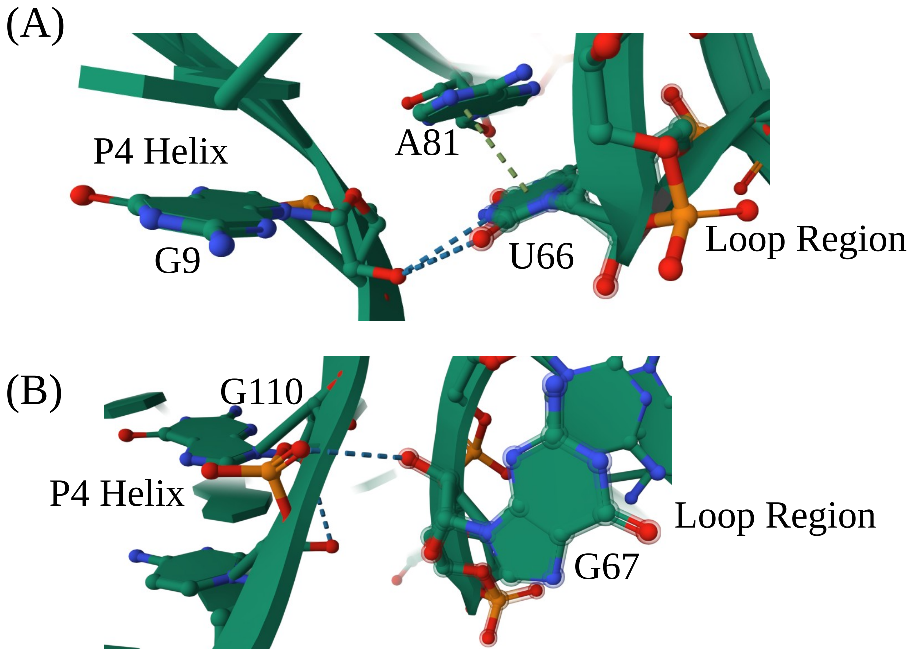


Figure S8: (A) Base-sugar non-canonical hydrogen bonds (blue dotted line) between G9 of P4 helix and U66 of loop region in the crystal structure. (B) Phosphate-sugar non-canonical hydrogen bonds (blue dotted line) between G110 of P4 helix and G67 of loop region in the crystal structure.

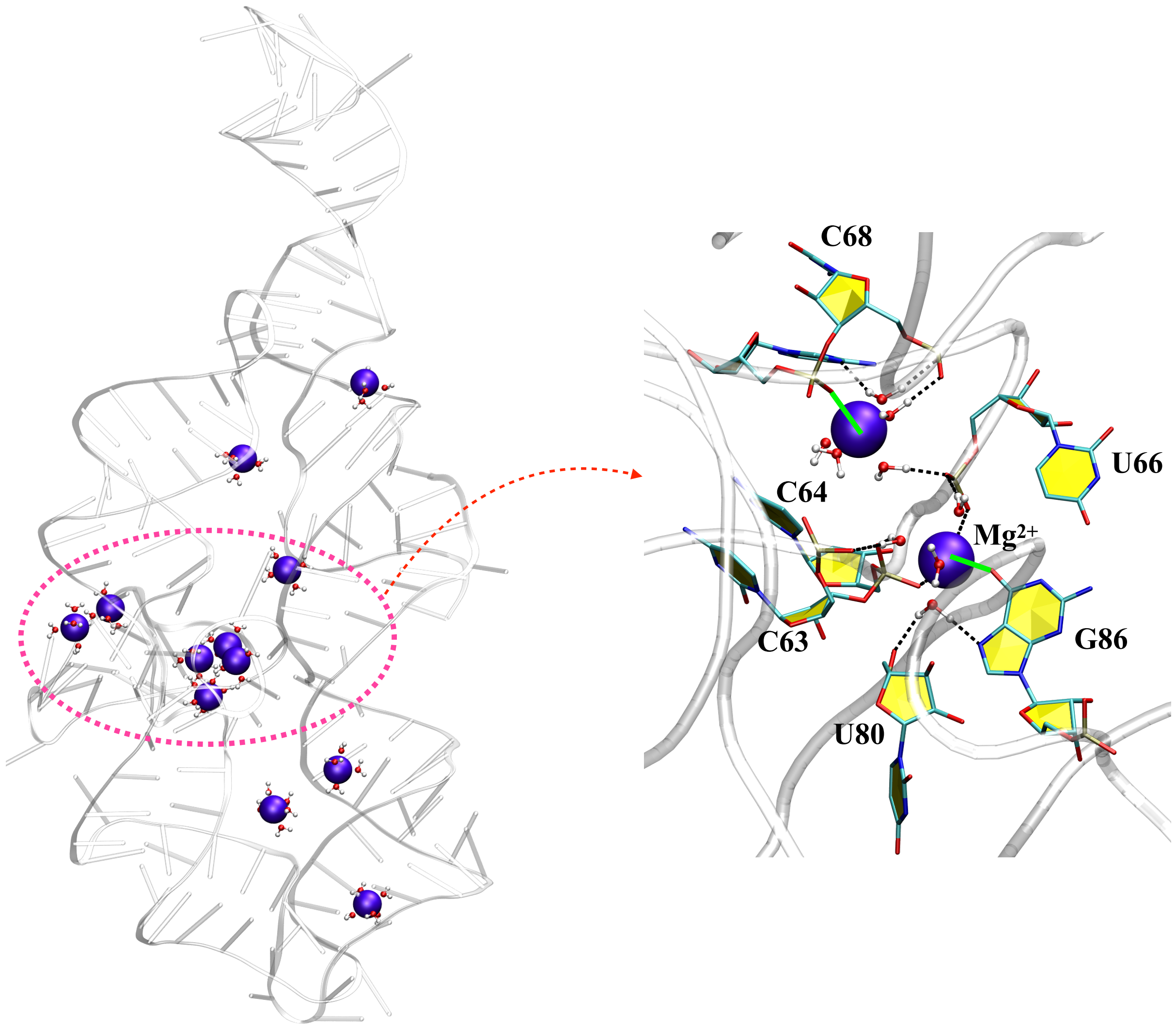


Figure S9: Crystal structure of P4-P6 domain (white) showing hydrated $Mg^{2+}$ ions (violet). $Mg^{2+}$ ions form IS (dotted line) and OS (green line) coordinations with RNA heavy atoms.

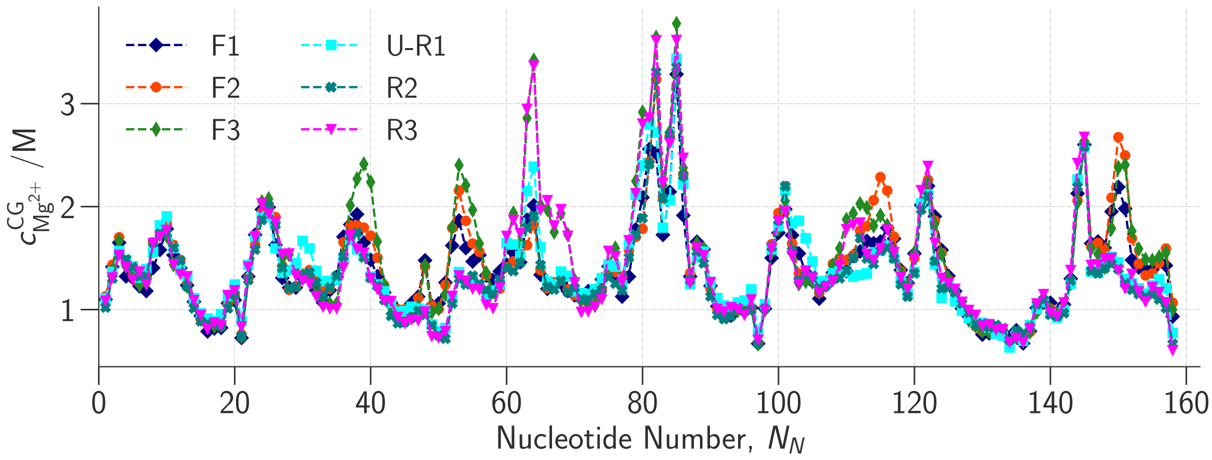


Figure S10: Local ion concentration for $Mg^{2+}$ ($c^{\mathrm{CG}}_{\mathrm{Mg}^{2+}}$) around the phosphate sites for six states (U-R1, R2, R3, F1, F2 and F3).

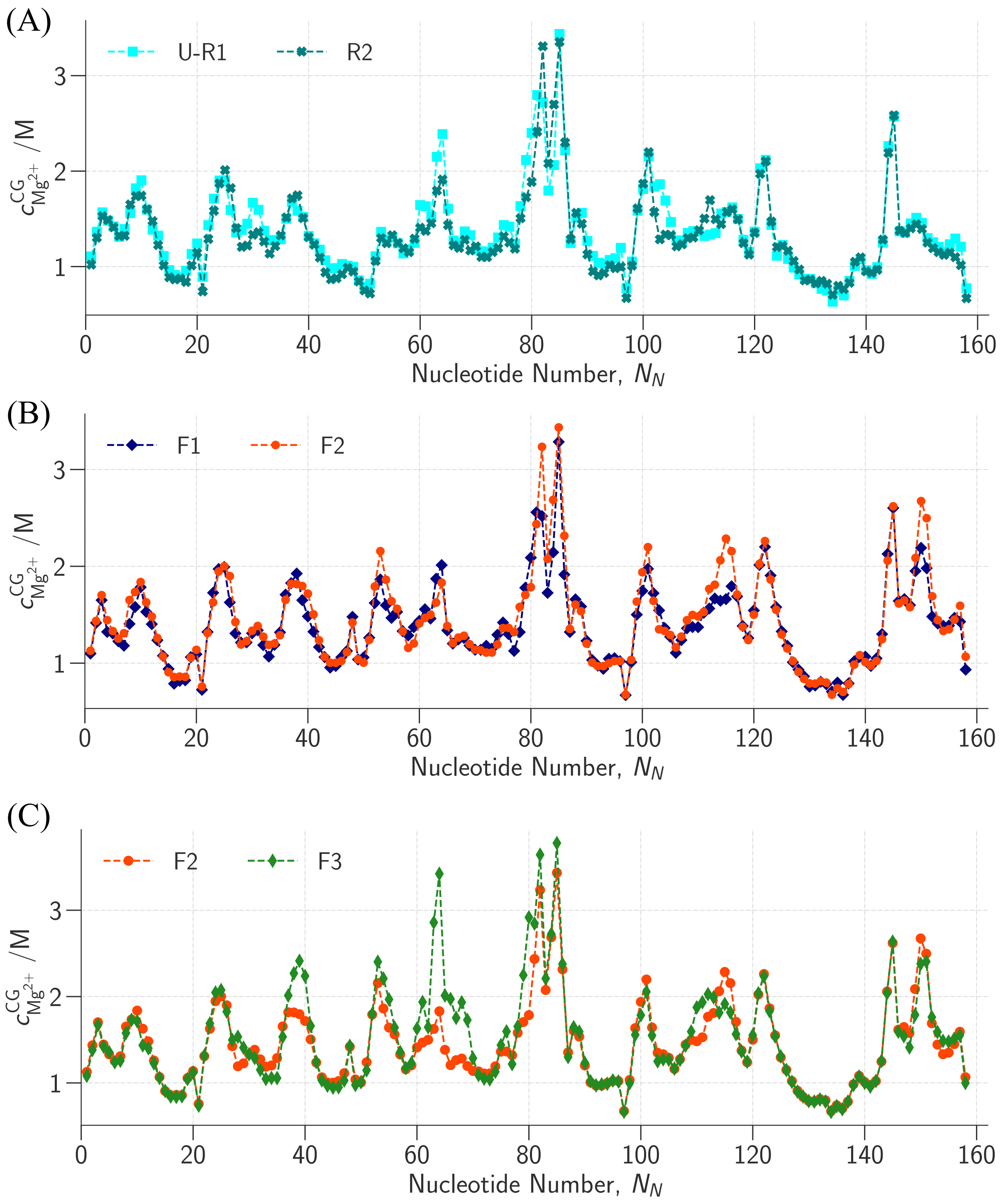


Figure S11: Local ion concentration for $Mg^{2+}$ ($c^{CG}_{Mg^{2+}}$) around the phosphate sites for (A) U-R1 and R2 states, (B) F1 and F2 states, (C) F2 and F3 states.

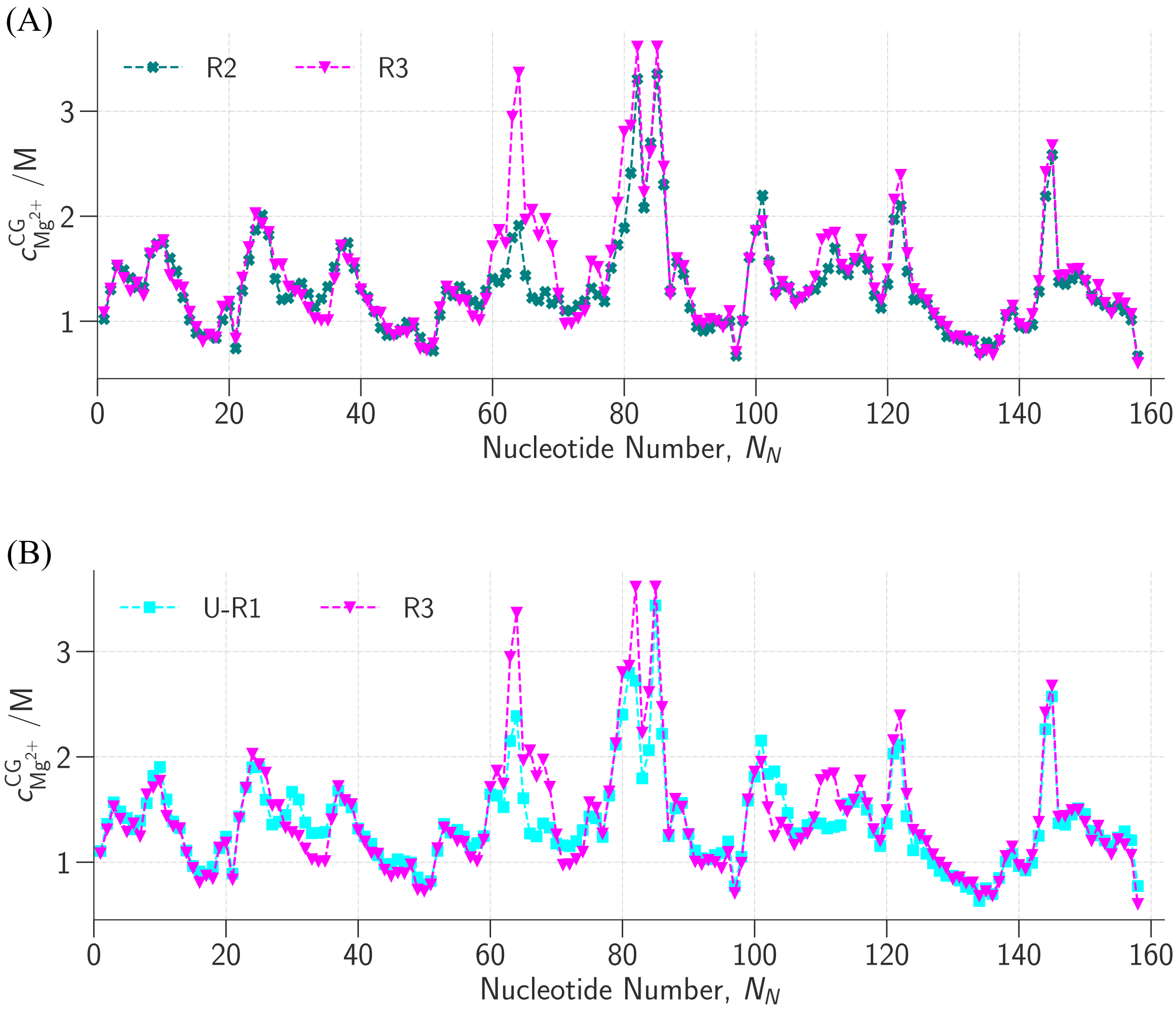


Figure S12: Local ion concentration for $Mg^{2+}$ ($c^{CG}_{Mg^{2+}}$) around the phosphate sites for (A) R2 and R3 states, (B) U-R1 and R3 states.

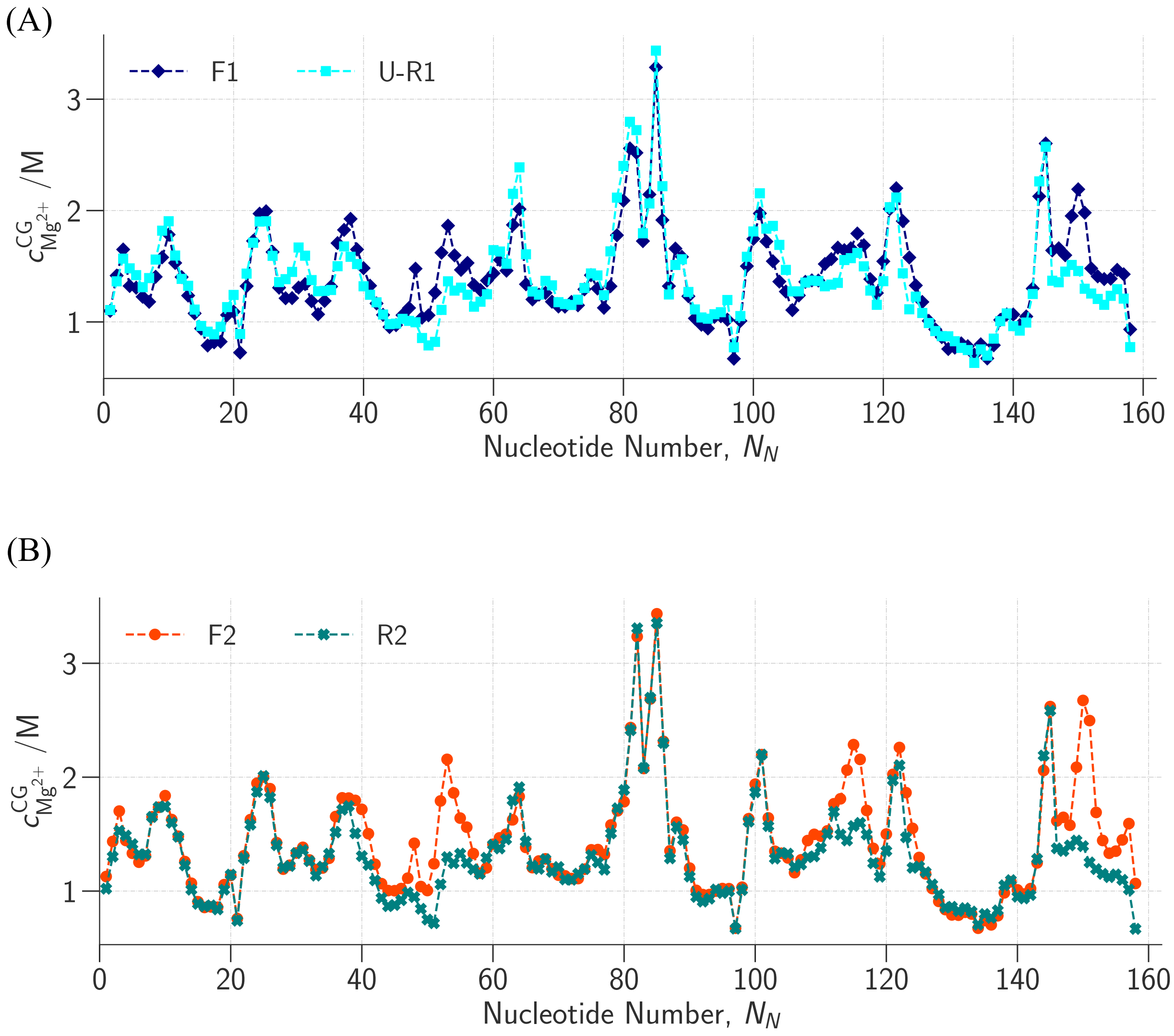


Figure S13: Local ion concentration for $Mg^{2+}$ ($c^{CG}_{Mg^{2+}}$) around the phosphate sites for (A) U-R1 and F1 states, (B) R2 and F2 states.

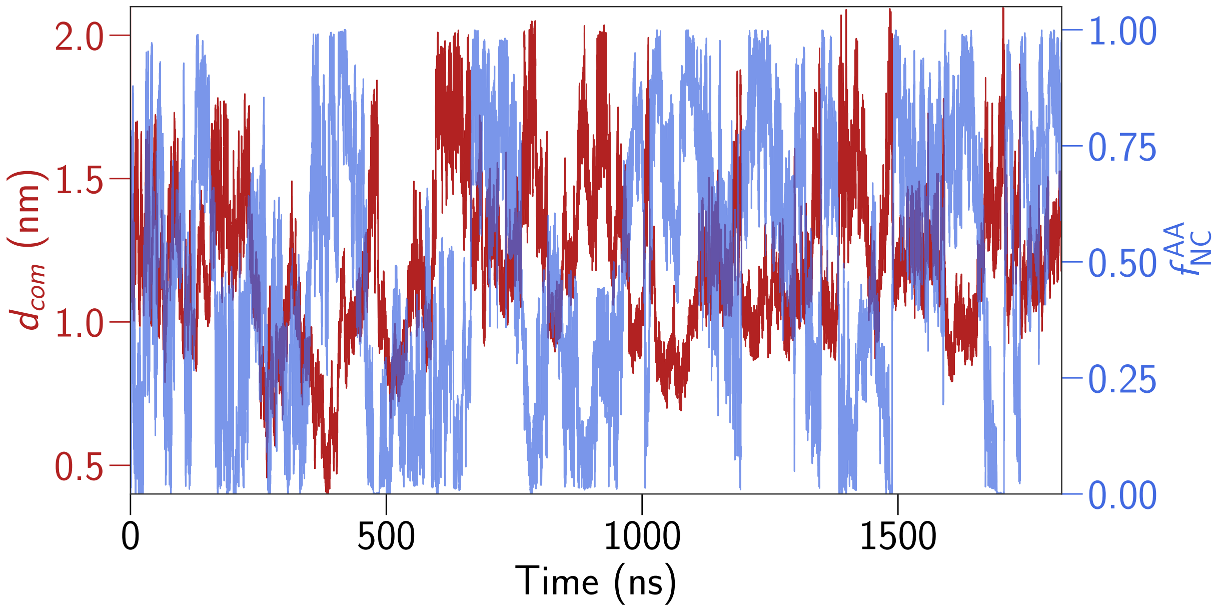


Figure S14: $d_{com}$ and $f_{\text{NC}}^{\text{AA}}$ plotted as a function of time in the WT-MetaD simulation trajectory.

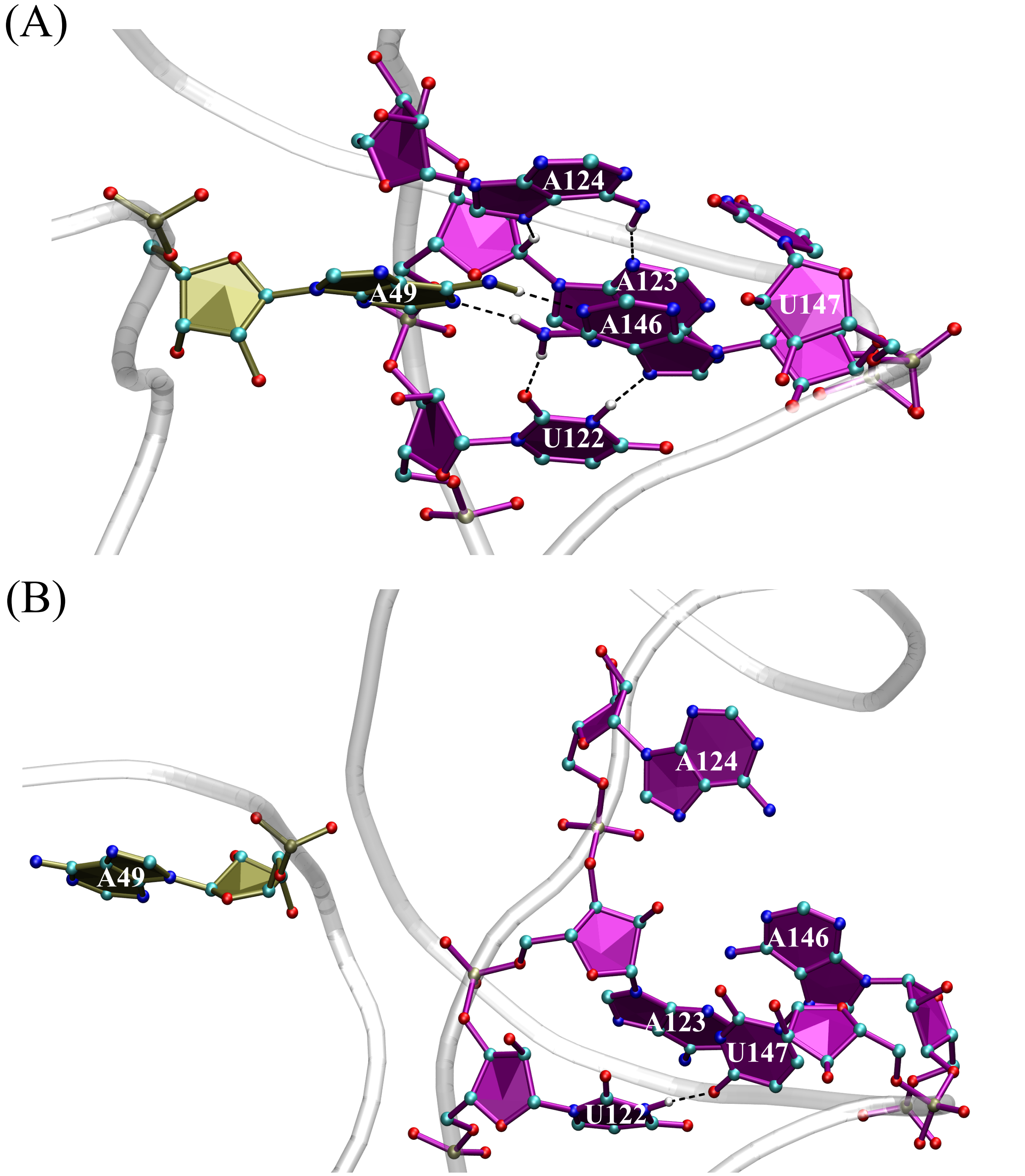


Figure S15: Hydrogen bonds between TL helix (A49) and R helix (U122, A123, A124, A146 and U147) in (A) TL-R docked state, and (B) TL-R undocked state.

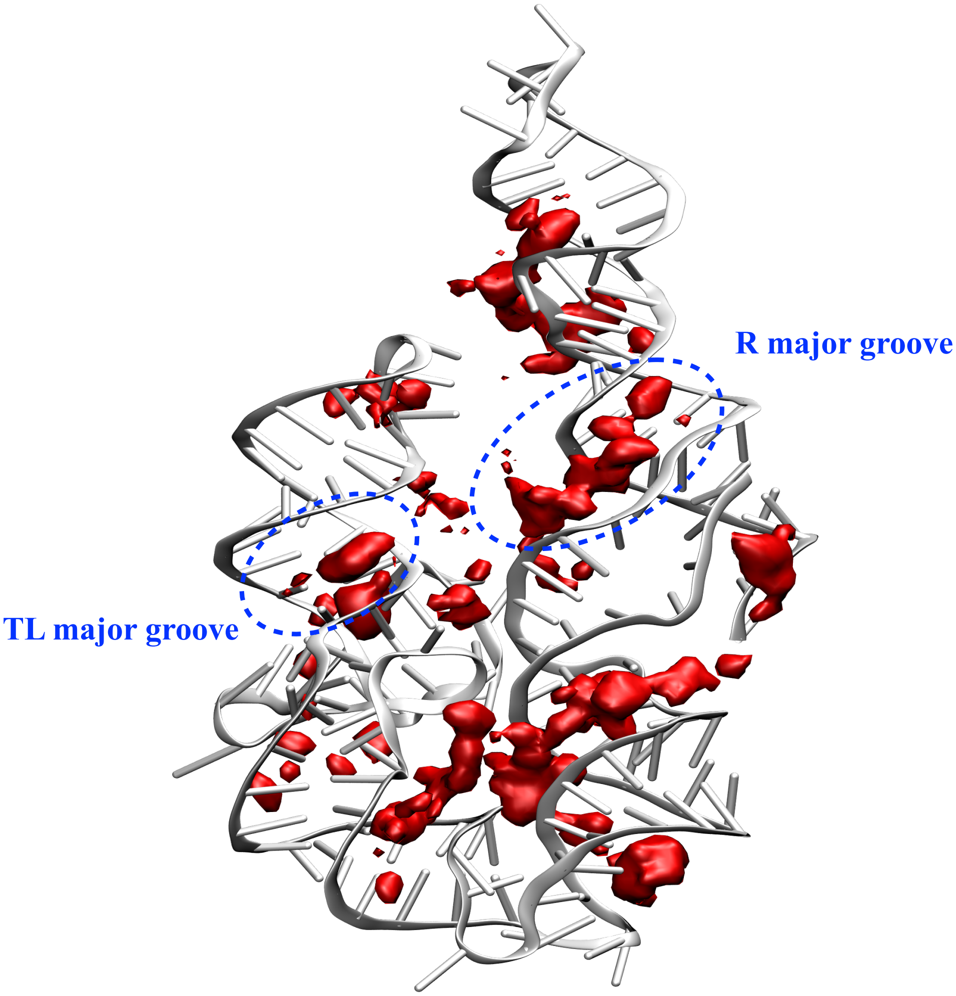


Figure S16: Spatial density map of $Mg^{2+}$ ions shown as red isosurfaces in the R3 state from all-atom simulations.

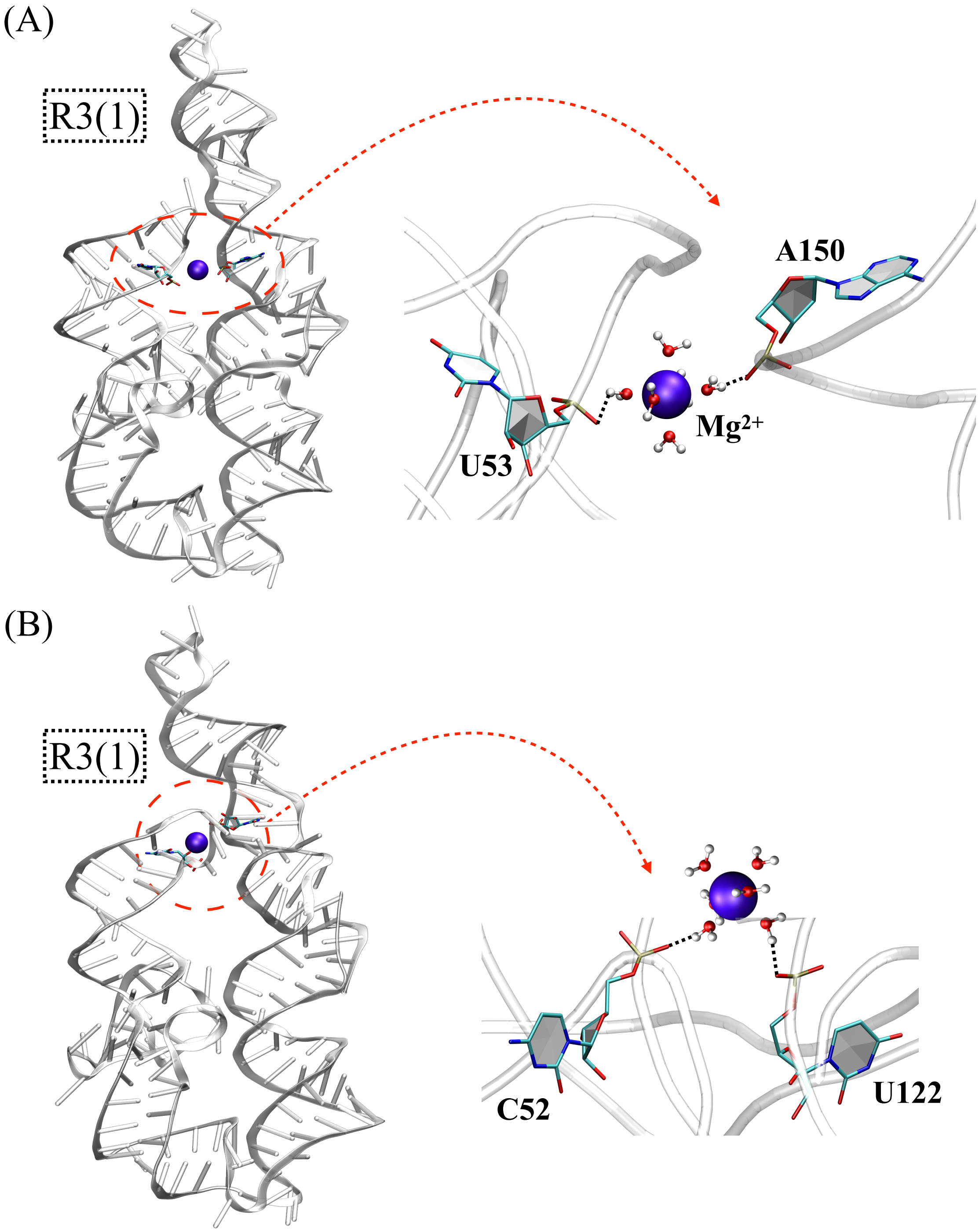


Figure S17: (A) and (B) In the R3(1) state, water mediated OS interactions (dotted black lines) between the $Mg^{2+}$ (violet coloured) ions and phosphate oxygens of nucleotides in the TL and R helices.

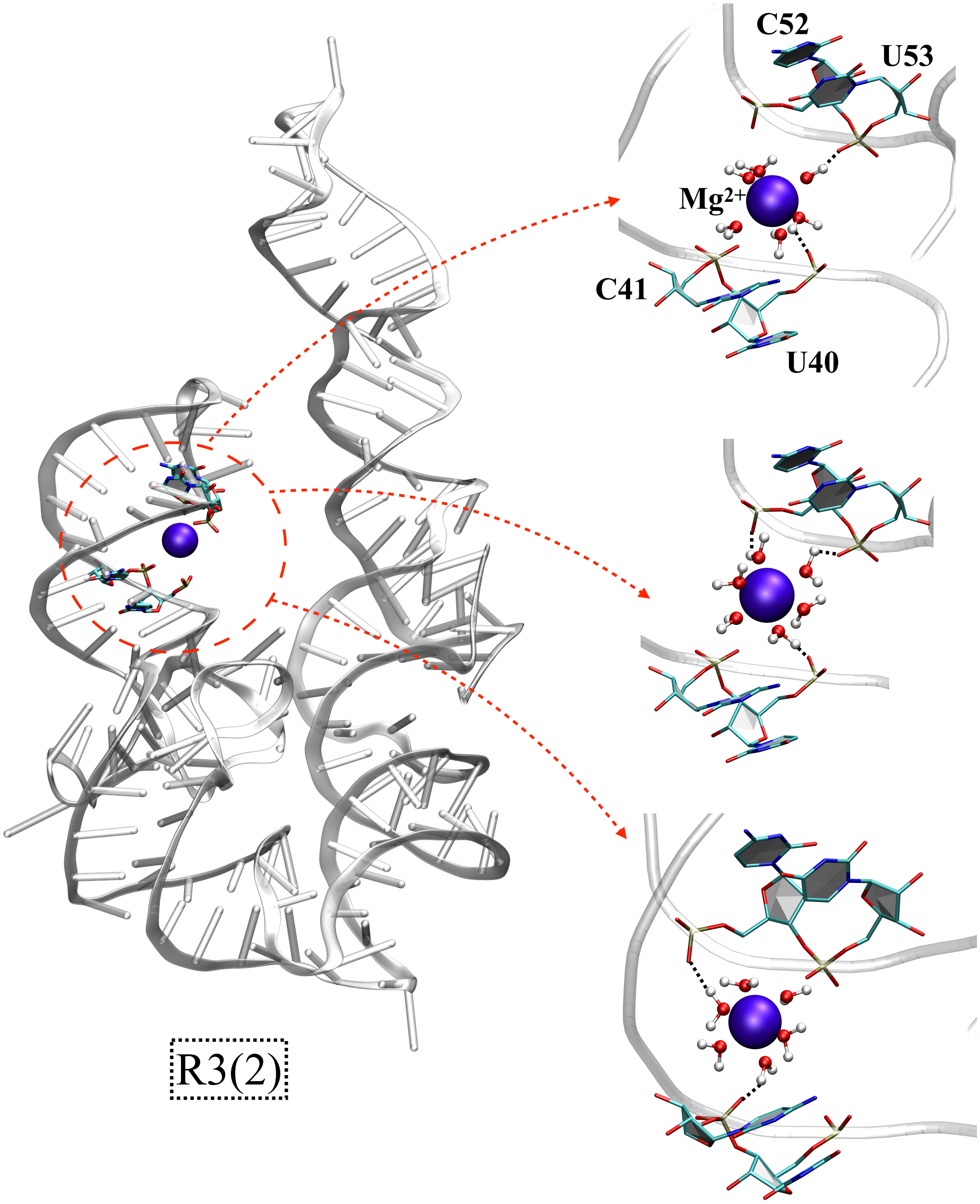


Figure S18: In the R3(2) state, water mediated OS interactions (dotted black lines) between the $Mg^{2+}$ (violet colored) ions and phosphate oxygens of nucleotides in TL helix.

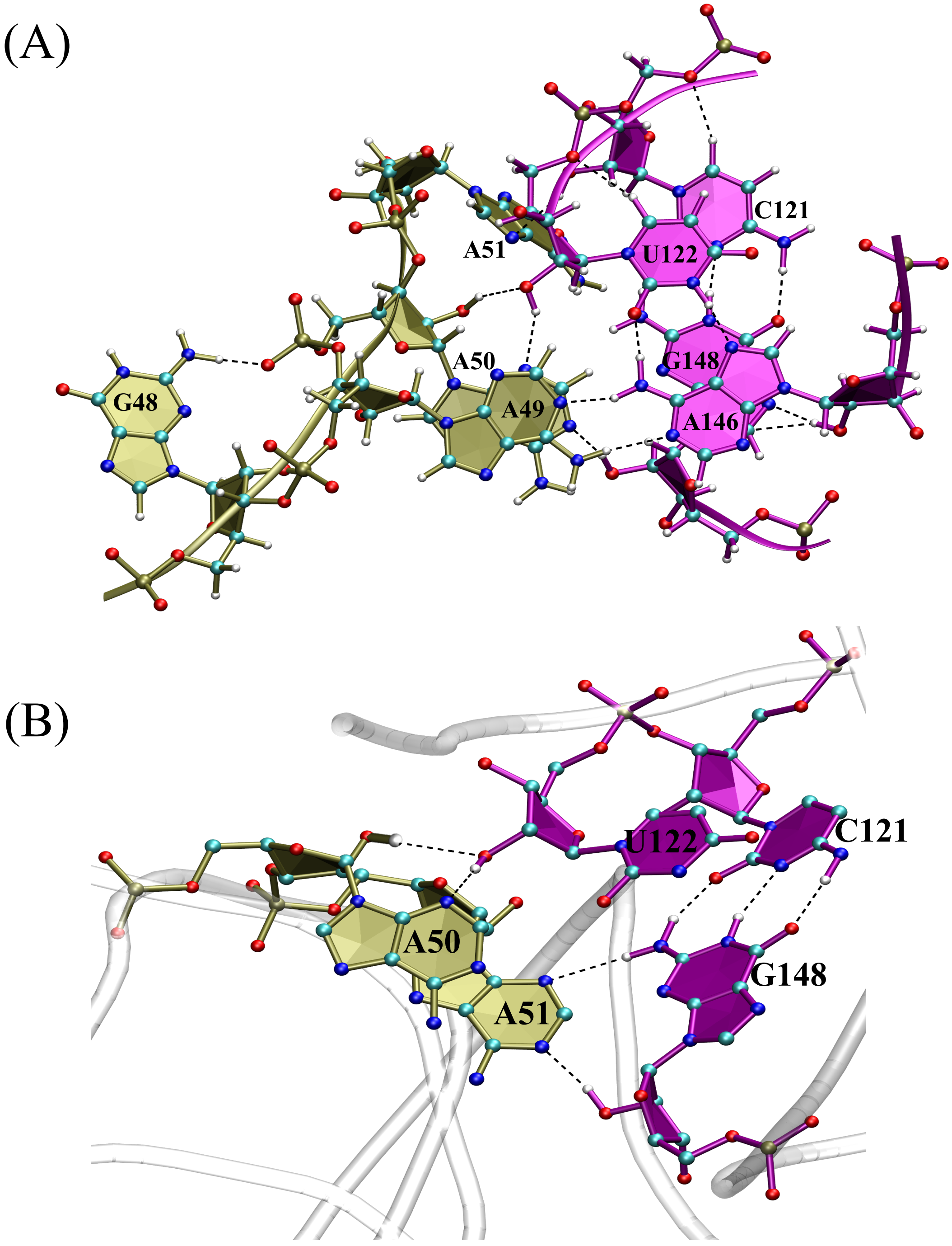


Figure S19: Native hydrogen bonds (dotted line) formed between atoms in TL-R docked complex in F3 state (A) in the presence of triplex H-bonds (A49-A146-U122) (B) in the absence of triplex H-bonds (A49-A146-U122).

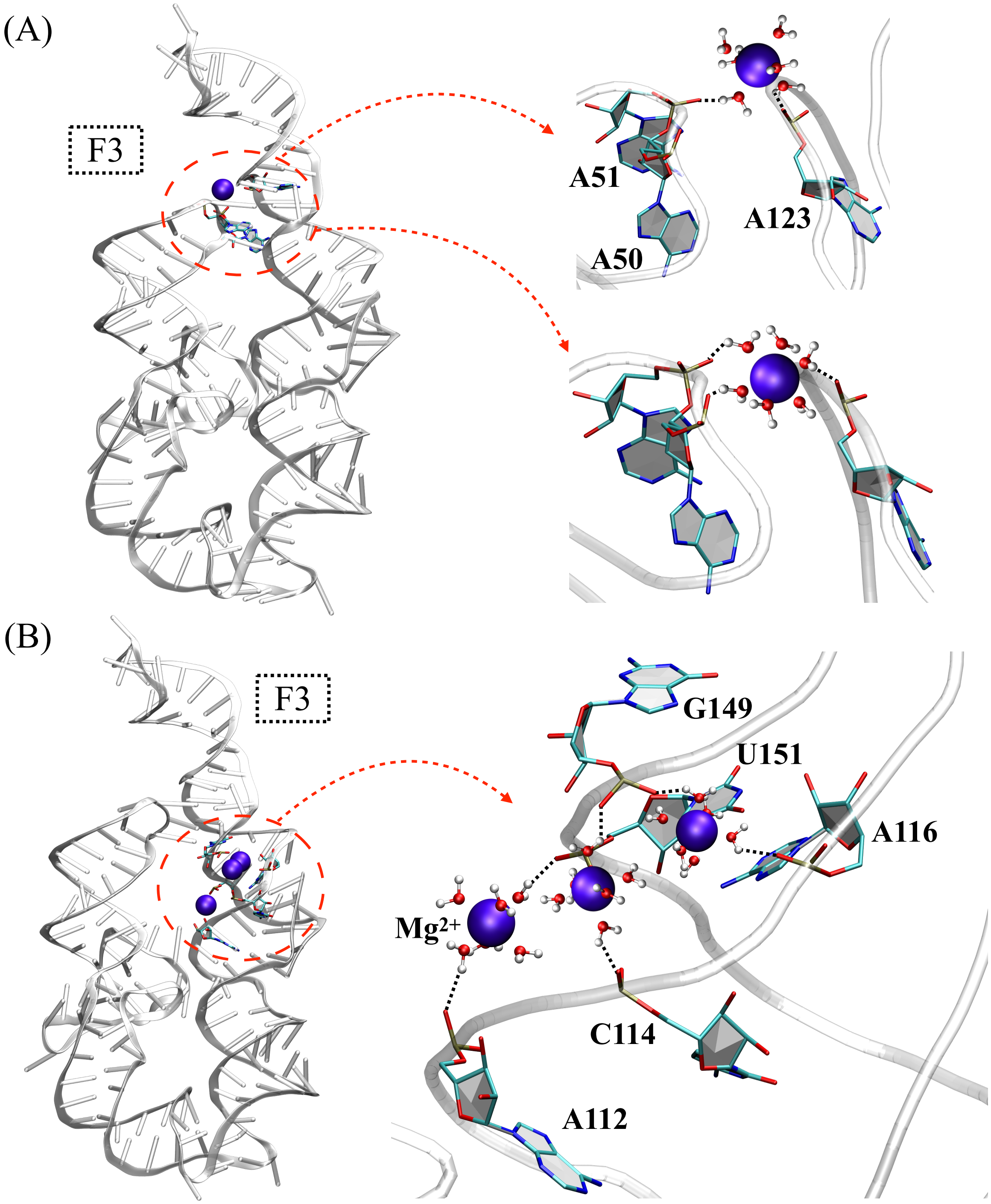


Figure S20: In the F3 state, water mediated OS interactions (dotted black lines) between the $Mg^{2+}$ (violet colored) ions and phosphate oxygens of nucleotides in the major groove of R helix.

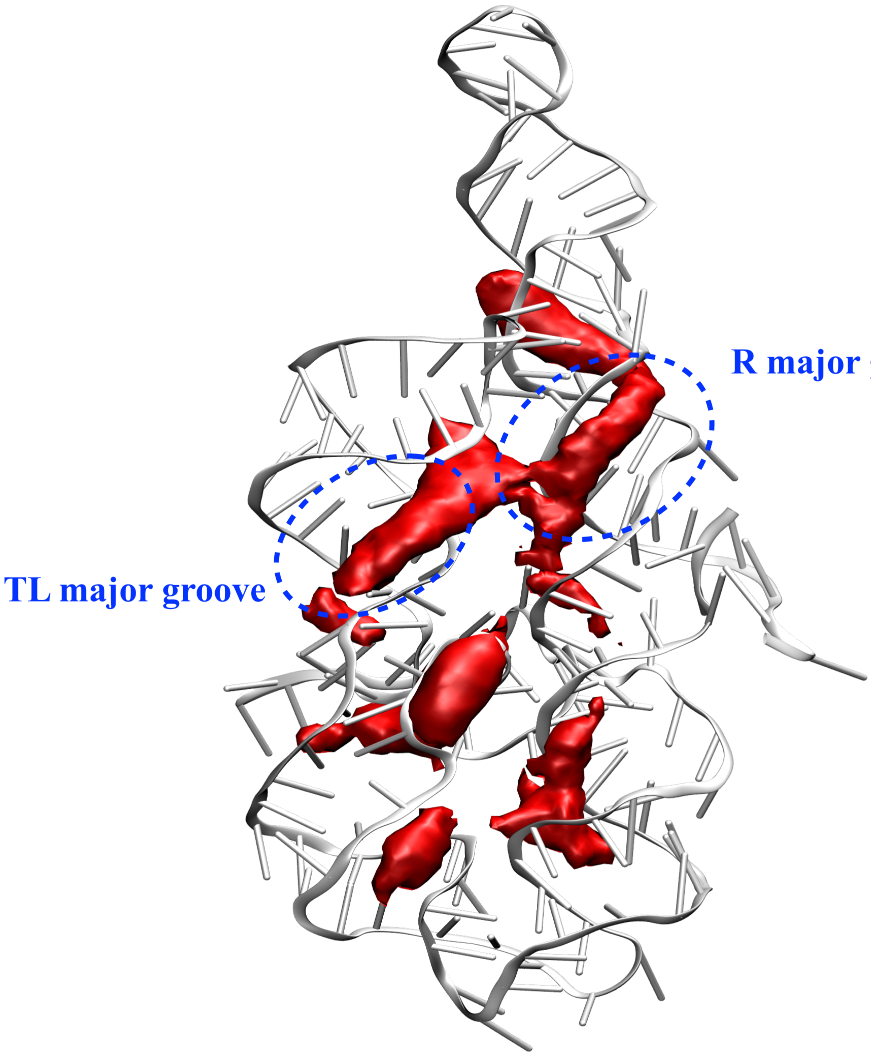


Figure S21: In the F3 state obtained from CG simulations, spatial density map of $Mg^{2+}$ ions condensed in the TL and R major grooves are shown as red isosurface.
